# A single-cell atlas of the *Culex tarsalis* midgut during West Nile virus infection

**DOI:** 10.1101/2024.07.23.603613

**Authors:** Emily A. Fitzmeyer, Taru S. Dutt, Silvain Pinaud, Barb Graham, Emily N. Gallichotte, Jessica L. Hill, Corey L. Campbell, Hunter Ogg, Virginia Howick, Mara K. N. Lawniczak, Erin Osborne Nishimura, Sarah Hélène Merkling, Marcela Henao-Tamayo, Gregory D. Ebel

**Affiliations:** Department of Microbiology, Immunology and Pathology, College of Veterinary Medicine and Biomedical Sciences, Colorado State University, Fort Collins, Colorado, USA; MIVEGEC, Université de Montpellier, IRD, CNRS, Montpellier, France; Department of Biochemistry and Molecular Biology, College of Natural Sciences, Colorado State University, Fort Collins, Colorado, USA; School of Biodiversity, One Health and Veterinary Medicine, Wellcome Centre for Integrative Parasitology, University of Glasgow, UK; Tree of Life, Wellcome Sanger Institute, Hinxton, CB10 1SA, UK; Institut Pasteur, Université Paris Cité, CNRS UMR2000, Insect-Virus Interactions Unit, 75015 Paris, France

## Abstract

The mosquito midgut functions as a key interface between pathogen and vector. However, studies of midgut physiology and virus infection dynamics are scarce, and in *Culex tarsalis* – an extremely efficient vector of West Nile virus (WNV) – nonexistent. We performed single-cell RNA sequencing on *Cx. tarsalis* midguts, defined multiple cell types, and determined whether specific cell types are more permissive to WNV infection. We identified 20 cell states comprising 8 distinct cell types, consistent with existing descriptions of *Drosophila* and *Aedes aegypti* midgut physiology. Most midgut cell populations were permissive to WNV infection. However, there were higher levels of WNV RNA (vRNA) in enteroendocrine cells, suggesting enhanced replication in this population. In contrast, proliferating intestinal stem cells (ISC) had the lowest levels of vRNA, a finding consistent with studies suggesting ISC proliferation in the midgut is involved in infection control. ISCs were also found to have a strong transcriptional response to WNV infection; genes involved in ribosome structure and biogenesis, and translation were significantly downregulated in WNV-infected ISC populations. Notably, we did not detect significant WNV-infection induced upregulation of canonical mosquito antiviral immune genes (e.g., *AGO2*, *R2D2*, etc.) at the whole-midgut level. Rather, we observed a significant positive correlation between immune gene expression levels and vRNA load in individual cells, suggesting that within midgut cells, high levels of vRNA may trigger antiviral responses. Our findings establish a *Cx. tarsalis* midgut cell atlas, and provide insight into midgut infection dynamics of WNV by characterizing cell-type specific enhancement/restriction of, and immune response to, infection at the single-cell level.

**Author Summary:** West Nile virus is the leading cause of mosquito-borne disease in N. America. *Cx. tarsalis* is a highly competent vector of WNV that plays a central role in the transmission and maintenance of WNV in nature. It is hypothesized that the permissibility of mosquito midgut cells contributes to the midgut infection barrier and thus impacts the ability of pathogens to establish infection in a mosquito. Additionally, it is postulated that the midgut is the most important organ with respect to determining vector competence. The recent publication of the full *Cx. tarsalis* genome, in conjunction with the growing body of work demonstrating the successful application of single-cell RNA sequencing methodologies in insect models made it possible for us to examine the cellular composition of the *Cx. tarsalis* midgut, and WNV infection dynamics therein, at single-cell resolution. We found cell-type-specific differences in viral RNA levels suggesting variability in WNV replication efficiency in specific cell types, identified patterns of differential expression associated with WNV infection in specific cell populations, and characterized aspects of the innate immune response to WNV infection at the tissue and cellular level.

## Introduction

Arthropod-borne viruses represent a severe and ever-growing public health threat (1). Mosquito-borne viruses alone are estimated to cause over 400 million infections globally each year (2). For transmission of a mosquito-borne virus to occur, a mosquito must first become infected with a virus via ingestion of an infectious bloodmeal after feeding on a viremic host (3,4). The virus must establish infection in the mosquito midgut before it disseminates into the hemocoel, and eventually enters the salivary glands and saliva – where transmission occurs (3,4). The mosquito midgut is a complex organ composed of a variety of cell types with distinct functions including digestion, nutrient absorption, endocrine signaling, and innate immune activity (5,6). The midgut is also the site of infection and escape barriers that strongly influence virus population dynamics (4,5). Previous studies have demonstrated that successful infection of the midgut epithelium, and replication and immune evasion therein, is essential for establishing disseminated infection in an arthropod vector (3,7). In these ways, for hematophagous disease vectors like mosquitoes, the midgut serves as a critical interface between vector and pathogen and the establishment of infection.

*Cx. tarsalis* is a major vector of West Nile virus (WNV) in much of North America (8–10). WNV is the most epidemiologically important arbovirus in N. America, causing ∼2,700 deaths from 1999 to 2022 (8,9,11). Despite the importance of *Cx. tarsalis* as a vector of WNV and other important human viruses, studies examining the cellular composition of its midgut, and WNV infection dynamics therein, are nonexistent. The recent publication of the full *Cx. tarsalis* genome, in conjunction with the growing body of work demonstrating the successful application of single-cell RNA sequencing methodologies in insect models has made it possible to address this significant knowledge gap (12–20). Therefore, we performed single-cell RNA sequencing (scRNA-seq) on dissociated midgut cells from both mock and WNV-infected *Cx. tarsalis* mosquitoes to gain a better understanding of how the midgut functions as the interface between vector and WNV.

We utilized a scRNA-seq approach previously demonstrated to be flavivirus RNA inclusive, which allowed us to detect WNV viral RNA (vRNA) in addition to host transcripts (21). Through this approach we identified distinct midgut populations corresponding with midgut cell types previously described in *Drosophila* and *Aedes aegypti* midguts – enterocyte (nutrient absorption cells), enteroendocrine (secretory cells), cardia (peritrophic matrix secreting cells), intestinal stem cell/enteroblast (undifferentiated progenitor cells), proliferating intestinal stem cell/enteroblast, visceral muscle cells, and hemocytes (immune cells) – and characterized the infection and replication dynamics of WNV within each population (5,6,14,16,17). We found that WNV infects most midgut cell types, with evidence suggesting enhanced replication in enteroendocrine cells and reduced viral replication in proliferating intestinal stem cells/enteroblasts. Additionally, we characterized the *Cx. tarsalis* immune response to WNV infection at both the whole-midgut and single-cell level. This study has bolstered our understanding of WNV midgut infection in a highly competent vector, and elucidated the midgut biology of *Cx. tarsalis*.

## Results

### Single-cell RNA sequencing of female *Cx. tarsalis* midguts identified 20 distinct cell populations

Using the 10X Genomics platform we performed scRNA-seq on pools of 10 dissociated *Cx. tarsalis* midguts at 4 and 12 days post-exposure to either an infectious bloodmeal containing WNV,or a mock uninfected bloodmeal.

Replicate infected and mock pools were collected and sequenced at each timepoint (four replicates at 4dpi and 2 replicates at 12dpi for each condition). We recovered an average of 2,416 cells per pool with an average coverage of 255,000 reads per cell, which were mapped to the *Cx. tarsalis* genome (**Supplemental File 1**).

Following quality control (QC) filtering, we retained data for 12,886 cells at 4dpi (7,386 WNV-infected, 5,500 mock), and 9,301 cells at 12dpi (4,609 WNV-infected, 4,692 mock) for downstream analyses (**Supplemental File 1**). Cells retained after QC contained an average of 597 (611 WNV-infected, 580 mock) and 407 (448 WNV-infected, 367 mock) unique genes per cell at days 4 and 12dpi respectively.

Guided clustering in Seurat (v4.3.0.1) generated 20 (4 dpi) and 17 (12 dpi) distinct clusters of cells (**Figure 1A-B**). The cell types of 15 cell clusters were identified using canonical gene markers and gene enrichment patterns previously identified in *Drosophila* and *Ae. aegypti* midguts (**Figure 1A-C**) (14,16–18,20). All cell-type identifications, with indicated exceptions, were based on cluster markers that were conserved between mock and WNV infection (**Supplemental File 2, 3**). We identified enterocytes (EC) by significant expression of *POU2F1* (*nubbin*), *PLA2G6* (*phospholipase A2*), and *AGBL5* (*zinc carboxypeptidase*), and enteroendocrine cells (EE) by expression of *PROX1* (*prospero*) (**Figure 2C, Supplemental Figure 3A**) (14,17,22,23). High expression of *Mlc2* (myosin light chain), *Mhc* (myosin heavy chain), and *ACTB* (*actin*), allowed us to identify visceral muscle cells (VM); VM-1, VM-2 (**Figure 1C**) (14,17,22,24). We identified a population of hemocytes (HC) based on expression of *NIMB2* and *SPARC* (**Figure 1C**) (16,18). EC-like cells (EC-like-1, EC-like-2, EC- like-3) were identified as such based on enrichment for several serine protease and alpha amylase genes (**Figure 1C, Supplemental File 2**) (14,17). One cardia population (cardia-1) was identified by enrichment for sugar transport and chitin-binding genes, as well as several serine protease genes, and a second cardia population (cardia-2) was identified by expression of C-type lysozyme and sugar transporter genes (**Figure 1C**) (14,17). Intestinal stem cells/enteroblasts (ISC/EB) were identified by visualizing *klumpfuss* (*klu*) expression specific to these clusters via violin plot (**Supplemental Figure 3B)**. One of the ISC/EB clusters was significantly enriched for *PCNA* – a marker for cell proliferation – and therefore named ISC/EB-prol to reflect this (**Figure 1C, Supplemental Figure 6**) (25,26). A cluster that shared identical conserved markers with cardia-1 and was also significantly enriched for *PCNA* was identified as cardia-prol (**Figure 1C**) (25,26). A cluster of fat body cells (FBC) was identified by high expression of *apoliphorin-III* (23). A cluster of Malpighian tubule cells (MT) that was only present in one out of 12 samples (mg5c) (**Supplemental Figure 2C**) was identified by significant enrichment for an inward rectifier potassium channel gene (*irk-2*) as well as several glutathione and vacuolar ATPase genes (**Figure 1C, Supplemental File 2**) (27,28). This indicates that Malpighian tubule tissue was inadvertently retained upon midgut collection for sample mg5c. Clusters without identifying markers are subsequently referred to by cluster number (e.g., cluster 4). Importantly, HCs, FBCs, and MTs are not midgut cells, but are considered associated with the midgut, while EC, EE, cardia, ISC/EB, and VM cell populations comprise the midgut, and EC, EE, cardia, and ISC/EB comprise the midgut epithelium (**Figure 2A-B**). We compared the proportion of each cluster between mock and WNV-infected replicates and found no significant differences (**Supplemental Figure 2A-B**). The percent of the total population composed by each cluster/cell-type can be found in **Supplemental Table 1**. Finally, we visualized the expression of S and G2/M phase cell cycle markers and found no evidence of cell cycle-driven clustering, with the exception of proliferating cell populations (**Supplemental Figures 8 & 9**).

**Figure 1.**
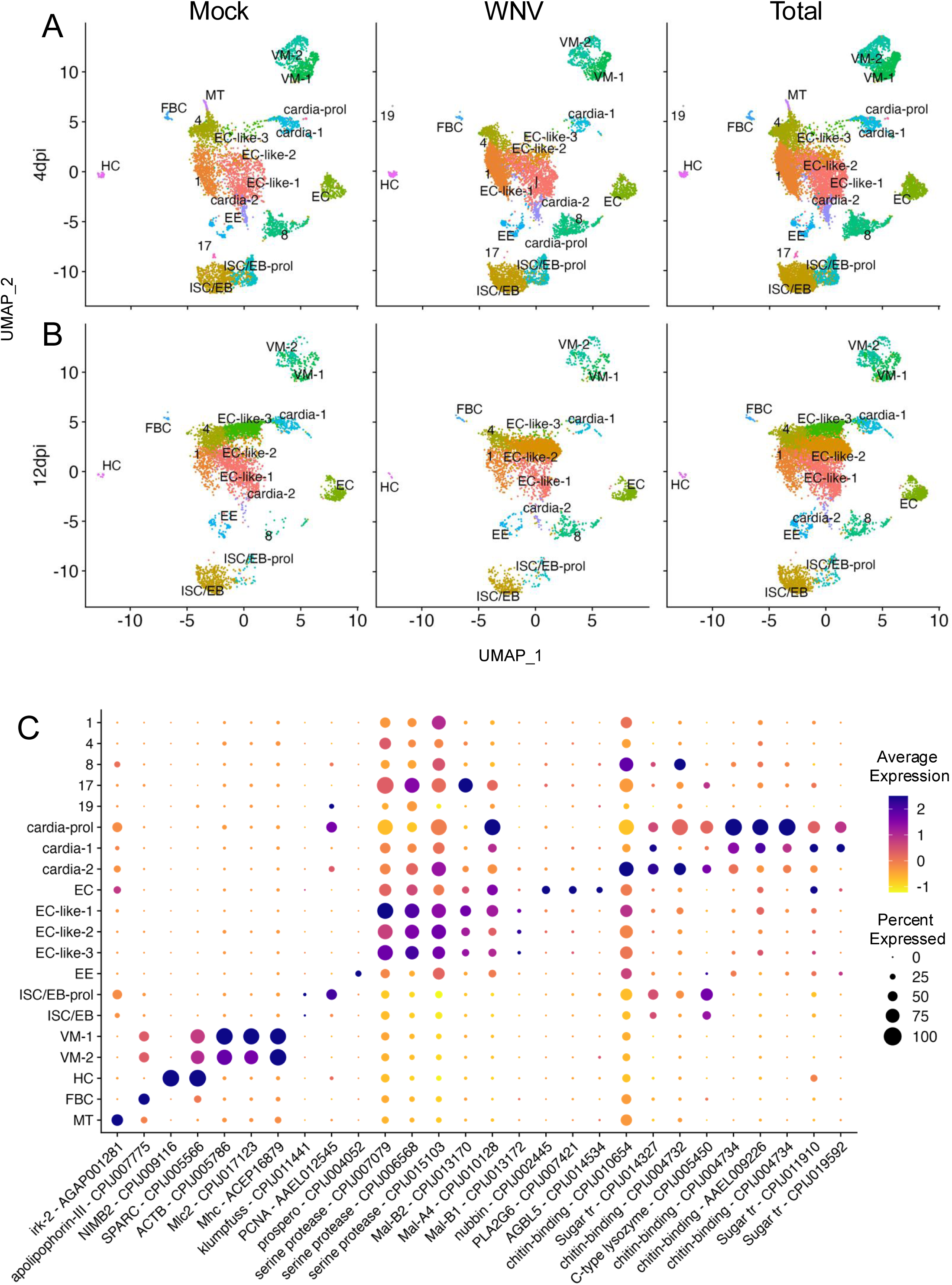
Single-cell sequencing of *Cx. tarsalis* midguts and cell-typing of midgut cell populations. Uniform Manifold Approximation and Projection (UMAP) reduction was used to visualize midgut cell populations. (**A**) 20 distinct clusters were identified at 4dpi and (**B**) 17 distinct clusters were identified at 12dpi. (**C**) Expression of canonical marker genes that were conserved between infection condition (mock and WNV-infected) were used to determine cell type. Dot size denotes percent of cells expressing each gene, color denotes scaled gene expression. Panel C was derived from the total population of cells for each subtype.

**Figure 2.**
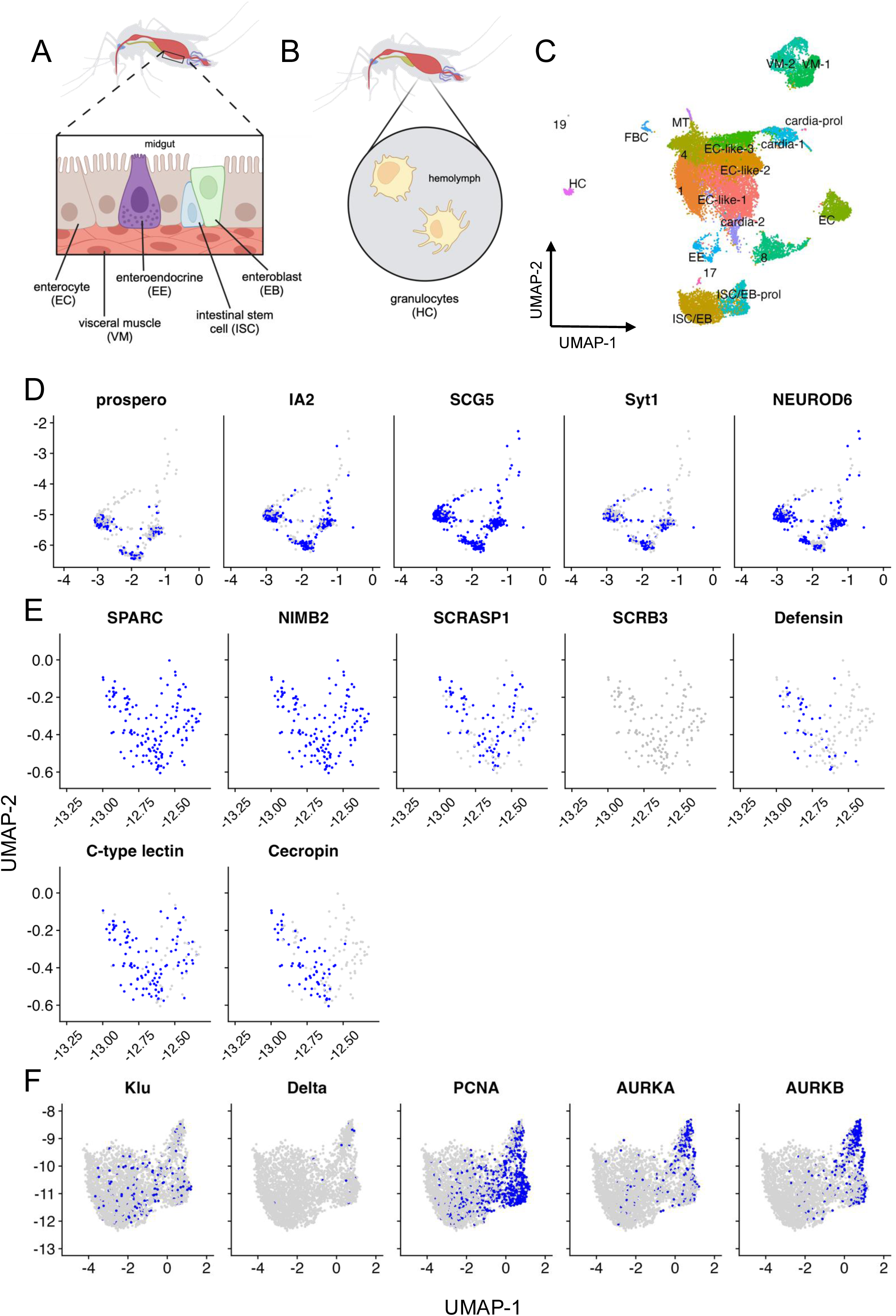
Further characterization of enteroendocrine cells, hemocytes and progenitor cells. Expression of characterizing genes visualized for EEs (**A**), HCs (**B**), and ISC/EBs (**A**) Gene counts for the total population (timepoints and conditions combined) were converted to binary (Blue = 1, Grey = 0) and visualized via UMAP. UMAP limits were set to include only clusters of interest, orientation of clusters in the total population can be found in panel **C**. (**D**) Expression of canonical enteroendocrine/neuroendocrine, neuropeptide, and secretory genes. (**E**) Expression of canonical HC, and granulocyte and oenocytoid genes (**F**) Expression of canonical ISC and EB genes as well as gene markers for proliferation and mitosis. Panels A and B were created in BioRender. Gallichotte, E. (2024) https://BioRender.com/f66n058.

### Characterization of *Cx. tarsalis* midgut secretory, immune, and progenitor cells

Enteroendocrine cells (EE) are the secretory cells of the midgut (**Figure 2A**) that, through the secretion of neuropeptides, regulate digestion and behavioral responses associated with feeding, satiety, stress, etc. (14,17,29). These cells, and the neuropeptides they secrete, have been previously characterized in *Ae. aegypti* and *Drosophila*, but never *Cx. tarsalis* (30,31). We identified *Cx. tarsalis* orthologs for previously described insect gut hormones found in EE cells - *short neuropeptide* (*sNPF*), *bursicon* (*Burs*), *ion transport peptide* (*ITP*), and *tachykinin* (*Tk*) receptor (14,17,20,30–32). However, these neuropeptides/neuropeptide receptors were found to have very low and nonspecific expression in the EE population (**Supplemental Figure 3A**). The EE population was significantly enriched for canonical neuroendocrine genes (*IA2*, and *SCG5*) (33,34) and *Syt1*, and showed low and nonspecific expression of *Syt4*, *Syt6*, *Syx1A* and *nSyb* (genes associated with vesicle docking and secretion) (**Figure 2D, Supplemental Figure 3A)** (14,35). Interestingly, the EE population showed strong enrichment for *NEUROD6*, an ortholog for neurogenic differentiation factor 4 in *Aedes aegypti* (AAEL004252) and for a neuronal differentiation gene known to be involved in behavioral reinforcement in mammals (**Figure 2D, Supplemental Figure 3A)** (36).

Hemocytes (HC) are immune cells that circulate in the hemolymph (**Figure 2B**) and play various roles in the mosquito immune response that are HC class-dependent (15,16,18,37,38). Much like EEs, hemocytes have not been characterized in *Cx. tarsalis*. The HC population was identified as mature granulocytes by expression of *SCRASP1*, *c-type lectin*, *defensin*, and *cecropin* genes (**Figure 2E, Supplemental Figure 3C**) (16). We saw no expression of *SCRB3* – an oenocytoid marker – in our HC population (**Figure 2E, Supplemental Figure 3C**) providing further support for the observation that the HCs detected were granulocytes (18).

Intestinal stem cells (ISCs) are the progenitor cells of the midgut that differentiate into either enterocyte progenitors (enteroblasts, EBs) or enteroendocrine progenitors (EEPs) (**Figure 2A**) (39,40). We identified a *Culex* ortholog for *Delta* (*Delta*-like protein – CPIJ019429), a canonical ISC marker in *Ae. aeygpti* and *Drosophila* midguts, however, it was not enriched in our ISC populations (**Figure 2F**). EBs and ISCs are often indistinguishable by UMAP, and thus we were able to use *klumpfuss* (*Klu*), an EB marker, to identify ISC populations (14,17,20). The ISC/EB-prol population was enriched for *aurora kinase A* and *B* (mitotic markers) as well as *PCNA* (proliferation marker) further confirming the proliferative state of this cluster (**Figure 2F**) (25,26,41,42).

### COG profiles demonstrate homogeneity between midgut cell populations despite differences in conserved markers

We next examined the transcriptional profiles of each cluster to understand the function of unidentified clusters and compare the transcriptomes of distinct cell populations. We used cluster of orthologous genes (COG) categories, and a two-pronged approach to visualization – COG profiles of all genes expressed in >75% of cells in a cluster (termed “base genes”) and COG profiles of all significant (p<0.05), conserved cluster markers with positive log_2_ fold-changes (log_2_FC) relative to the other clusters (**Supplemental Figure 1A, B**). Base gene and cluster marker gene profiles were derived from the total population at each timepoint. Despite the varying cell types, we noted homogeneity across base genes for each cluster, with the plurality of each profile for most clusters comprising genes involved in translation and ribosomal biogenesis (J), and energy production and conversion (C) (**Supplemental Figure 1A**). However, ISC/EB-prol, cardia-2, and cardia-prol all possessed fewer ‘J’ COGs than other clusters. The COG profiles of EC-like populations show variability between their transcriptomes and ECs (**Supplemental Figure 1A-B**). As expected, VM populations contained the highest proportions of cytoskeletal genes (Z) in both base gene and cluster marker profiles compared to other cell types (**Supplemental Figure 1A-B**).

### WNV vRNA is detected at varying levels in the majority of midgut cell populations

In addition to characterizing the cellular heterogeneity of *Cx. tarsalis* midguts, we also sought to examine WNV infection dynamics at the single-cell level. The five-prime bias of the scRNA-seq chemistry captured and allowed us to detect the WNV 5’ UTR as a feature in our data. Visualization of the sequence alignment map files generated by the Cell Ranger pipeline confirmed that WNV reads mapped with Cell Ranger solely mapped to the 5’ UTR. Importantly, WNV viral RNA (vRNA) was only detected in our WNV-infected samples, and was broadly detected across most cell populations at both time points (**Figure 3A**). Within WNV-infected replicates we compared the percent of cells with detectable vRNA (calculated as percent expressing) and the average vRNA level (calculated as average expression) in the total population for each timepoint (**Figure 3B**). We saw no significant difference in the total percent of WNV-infected cells between timepoints, but a significant increase in average total vRNA level by 12dpi (**Figure 3B**). Within clusters (replicates within time points combined), cells contained variable levels of vRNA, however some clusters (cluster 17, cardia-prol, etc.) were either not present ≤5 cells in the WNV-infected condition (**Figure 3C**). Examining cluster markers in the WNV-infected condition revealed that vRNA was a significant cluster marker for several clusters at each timepoint (**Figure 3C**). At both timepoints, cluster 4 contained both the highest average level of vRNA and was significantly enriched for vRNA relative to the other clusters (**Figure 3C, Supplemental Figure 4, Supplemental Table 2**). Cluster 4 lacked canonical markers; however, it was significantly enriched for mitochondrial genes and mitochondrial tRNAs, suggesting these cells are either apoptotic or in states of increased energy demand or stress (**Supplemental Figure 4**). However, there was minimal expression of pro- apoptotic genes across all clusters, and no notable enrichment of pro-apoptotic genes in cluster 4 (**Supplemental Figure 7**). At both timepoints the ISC/EB-prol populations, and at 12dpi the ISC/EB population, were distinguished by a significant absence of vRNA relative to the other clusters (**Figure 3C**, **Supplemental Table 2**). vRNA in the 4dpi ISC/EB-prol population had a log_2_FC value of -2.2, and vRNA in the 12dpi ISC/EB- prol and ISC/EB populations had log_2_FC values of -3.6 and -2.5 respectively; greater absolute values than any other cell population/cluster (**Supplemental Table 2**).

**Figure 3.**
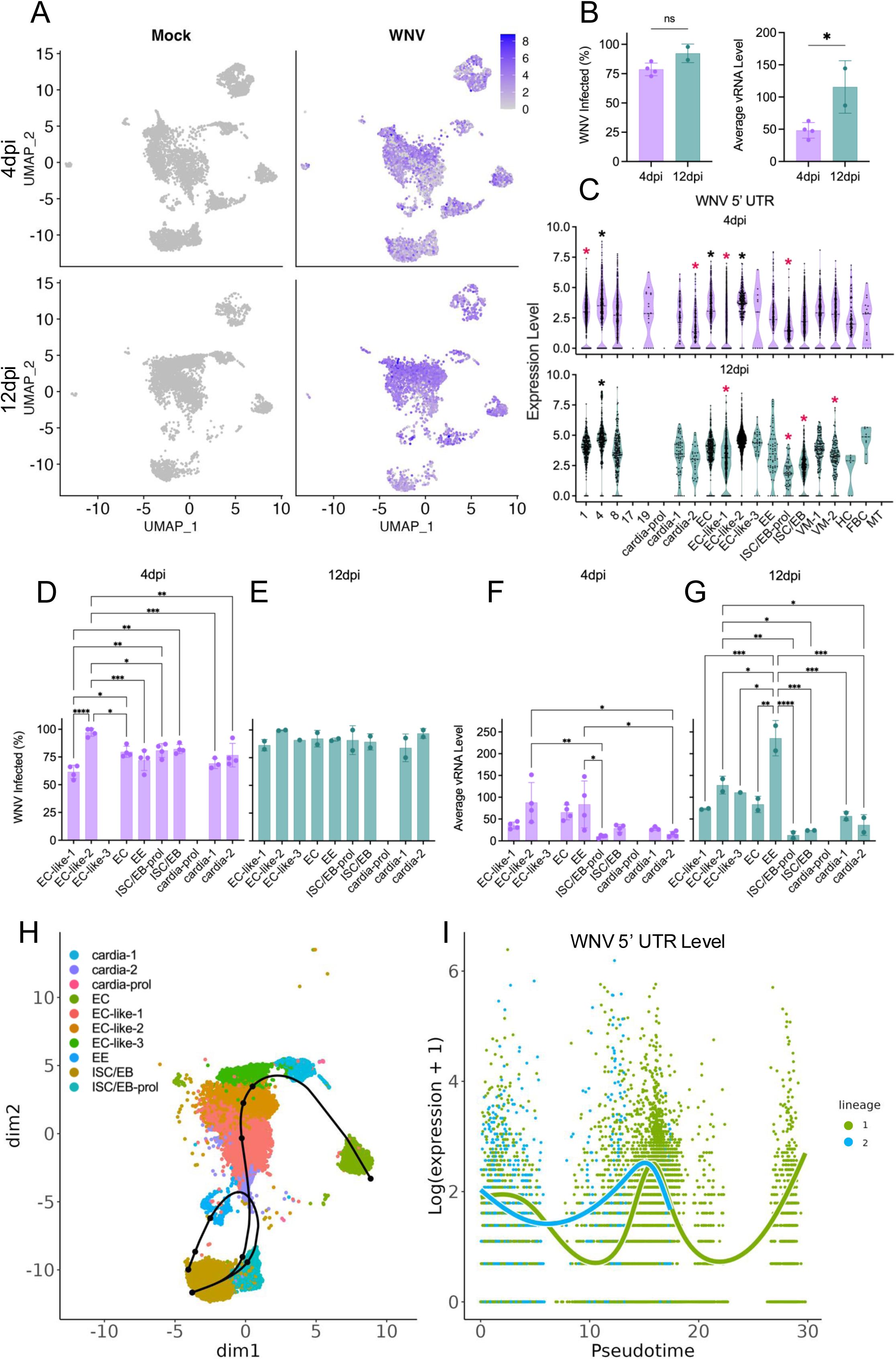
WNV vRNA is detected in most midgut cell populations. (**A**) Levels of WNV vRNA across midgut cell populations at 4 and 12dpi. Purple to grey gradient denotes high to low levels of vRNA. (**B**) percent of cells in the total population containing WNV vRNA (percent expressing) and average WNV vRNA level in the total population (average expression) calculated for each WNV-infected replicate and compared between timepoints. Significance was determined by unpaired t-test, * = p < 0.05. (**C**) WNV vRNA level in individual cells of distinct clusters. Black asterisks denote significant enrichment of vRNA as a cluster marker, pink asterisks denote significant absence of vRNA as a cluster marker. **Examining WNV vRNA levels in epithelial cell populations.** The percentage of cells containing WNV vRNA in each epithelial cell population at (**D**) 4dpi and (**E**) 12dpi. Average WNV vRNA level in epithelial cell populations at (**F**) 4dpi and (**G**) 12dpi. Significance determined by one-way ANOVA, * = p < 0.05, ** = p < 0.005, *** = p < 0.0005, **** = p < 0.0001. Points = replicate values, bars = mean of replicate values, error bars = SD. (**E**) Trajectory inference for ISC/EB and ISC/EB-prol populations identified two lineages. (**F**) vRNA levels visualized along lineage progression. Green = lineage 1 (EC), blue = lineage 2 (EE). Panels B-G were derived from only WNV-infected samples. Panels D-G exclude replicate values derived from clusters of ≤5 cells. Panels A, and H-I were derived from the total population.

### WNV vRNA levels differ between epithelial cell populations

Next, we sought to compare the presence and level of vRNA in epithelial cell populations; EC and EC-like, EE, cardia, ISC/EB, and ISC/EB-prol. Average expression and percent expression values derived from clusters composed of ≤ excluded from this comparison. At 4dpi the EC-like-2 population had the highest percentage of cells containing vRNA, significantly more than EC-like-1, EC, EE, ISC/EB-prol, and cardia populations (**Figure 3D**).

Interestingly, the other EC-like population at 4dpi (EC-like-1) had significantly lower percentages of cells containing vRNA compared to other populations (**Figure 3D**). There were no significant differences in the percent of cells containing vRNA between any epithelial cell population at 12dpi (**Figure 3E**). At both time points, EC-like-2 and EE populations the highest average levels of vRNA (**Figure 3F-G**). At 12dpi EE populations had significantly higher levels of vRNA than all other epithelial cell populations (**Figure 3G**).

To further explore the dynamics of epithelial cell populations we used slingshot (v2.10.0) to perform a trajectory inference and identify cell lineages. We identified two lineages: (1) ISC/EB → ISB/EB-prol → EE, and (2) ISC/EB → ISC/EB-prol → EC-like → EC (**Figure 3H**). As expected, we observed decreases in Klu and PCNA expression in both lineages with increasing pseudotime, and saw an increase in expression of EC cell marker nubbin and EE cell marker prospero corresponding with the differentiation of lineages 1 and 2 respectively (**Supplemental Figure 11**). Plotting levels of WNV vRNA along the progression of each lineage confirmed our finding that vRNA levels are lowest in the ISC/EB-prol population compared to fully differentiated EE and EC cells (**Figure 3I**). Importantly, visualizing vRNA along a lineage progression is a useful way to further examine vRNA level variability between cell populations, it is not a timeline of viral infection, i.e., it does not suggest that WNV initiates infection in the ISC/EB or ISC/EB-prol population.

### WNV infection is associated with downregulation of ribosomal and translation related genes in progenitor cells

To further examine the impact of WNV infection on specific cell populations, we performed pseudo-bulk differential expression (DE) analysis between mock and WNV-infected conditions for all midgut cell populations (EC, EC-like, EE, ISC/EB, cardia, VM). We identified 111 significant infection-associated DEGs in these populations with ISC/EB, ISC/EB-prol, and EC-like-1 populations comprising the greatest percentage of the total with 31, 25, and 24 DEGs respectively (**Figure 4A, Supplemental Table 3**). The majority (104/111) of cell-type specific, infection-associated DEGs were identified in populations from 4dpi, while only 7 DEGs were identified in populations from 12dpi (**Figure 4B**). The majority (89/111) of DEGs were downregulated in response to infection, with only 22 upregulated (**Figure 4B**). Interestingly, predominantly ribosomal and translation related genes were downregulated in infected ISC/EB and ISC/EB-prol populations at 4dpi (**Figure 4C, Supplemental Table 3**), and two innate immune genes were significantly downregulated in infected EE populations at 4dpi (**Figure 4C, Supplemental Table 3**). To get a broader overview of the biological processes and molecular functions associated with WNV infection, we performed gene ontology (GO) enrichment analysis of all DEGs associated with infection (**Figure 4D,E, Supplemental Table 3**). GO categories for genes with functions related to ribosomal structure and translation were enriched among DEGs downregulated in WNV-infected cell populations – likely driven by the large number of such genes downregulated in ISC/EB populations (**Figure 4D**). GO categories associated with amino acid salvage, and biological processes involving L-methionine were enriched among DEGs upregulated in WNV-infected cell populations (**Figure 4E**). However, due to there being fewer upregulated DEGs, the false discovery rates were much larger for the results of this GO enrichment analysis (**Figure 4E**).

**Figure 4.**
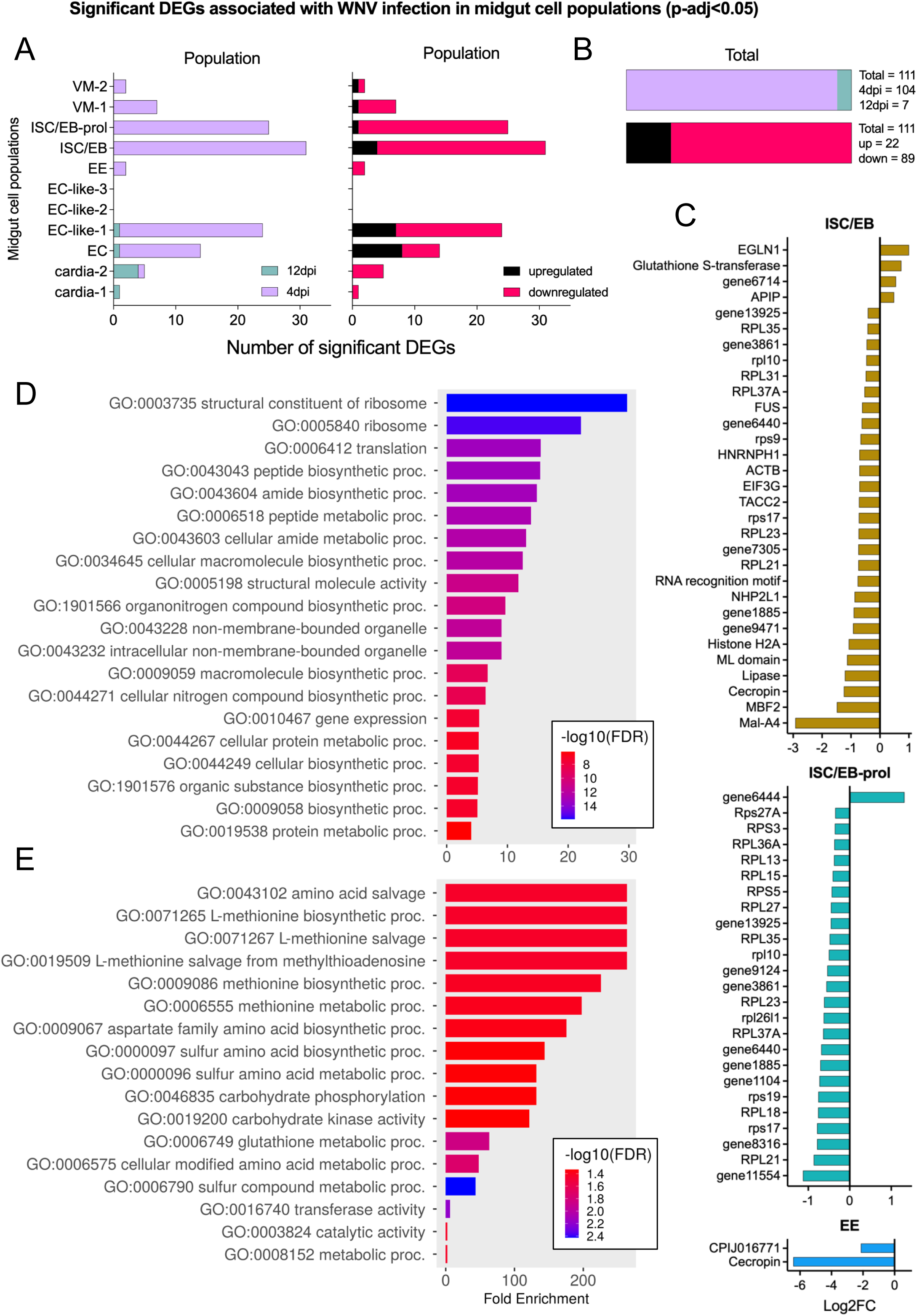
The impact of WNV infection on midgut cell populations. (**A**) Number of significant (adjusted p-value < 0.05) differentially expressed genes (DEGs) associated with WNV infection in each midgut cell type. Green = 12dpi, purple = 4dpi, black = upregulated, pink = downregulated. (**B**) Total significant DEGs identified in midgut cell populations. Green = 12dpi, purple = 4dpi, black = upregulated, pink = downregulated. (**C**) Visualizing WNV-infection associated DEGs in cell populations of interest (ISC/EBs, EEs). Negative log_2_FC indicates downregulation and positive log_2_FC indicates upregulation associated with WNV infection. In the interest of data density, the bar representing the log_2_FC of the WNV 5’ UTR is not shown. WNV 5’ UTR log_2_FC values can be found in **Supplemental Table 4**. (**D**) Gene ontology enrichment analysis (GOEA) for all downregulated DEGs (timepoints and cell populations combined). (**E**) GOEA for all upregulated DEGs (timepoints and cell populations combined). Color denotes -log10(FDR). FDR cutoff < 0.05 was used for GOEA.

### Identification of genes associated with WNV infection at the whole-tissue, and single-cell level

Bulk- RNA sequencing comparing WNV-infected to uninfected *Cx. tarsalis* midguts has not yet been described, so we performed a pseudo-bulk DE analysis to identify differentially expressed genes (DEGs) associated with mock and WNV-infected midguts at the whole-tissue level. We identified six significant DEGs at 4dpi; *homocysteine S-methyltransferase*, *DMAS1* (aldo-keto reductase), and *GBE1* (deltamethrin resistance-associated gene) were upregulated in response to WNV infection while *BCAN* (c-type lectin), uncharacterized gene11056, and a serine protease gene were downregulated (**Figure 5A, Supplemental Table 4.1**). At 12dpi we identified 10 significant DEGs; an ML (MD-2 related lipid recognition) domain-containing gene, four *CRYAB* (HSP20 heat shock protein) genes, and a chitin-binding domain-containing gene were upregulated in response to WNV infection, and three uncharacterized genes (gene13447, gene11056, gene9296) and fibrinogen/fibronectin were downregulated (**Figure 5B, Supplemental Table 4.1**). DEGs differed for each timepoint and, as such, we next examined DEGs in the WNV-infected condition between timepoints. We found many significant DEGs between timepoints; several leucine rich repeat containing genes were upregulated at 4dpi, and host immune gene *LYSC4* was upregulated at 12dpi (**Figure 5C**).

**Figure 5.**
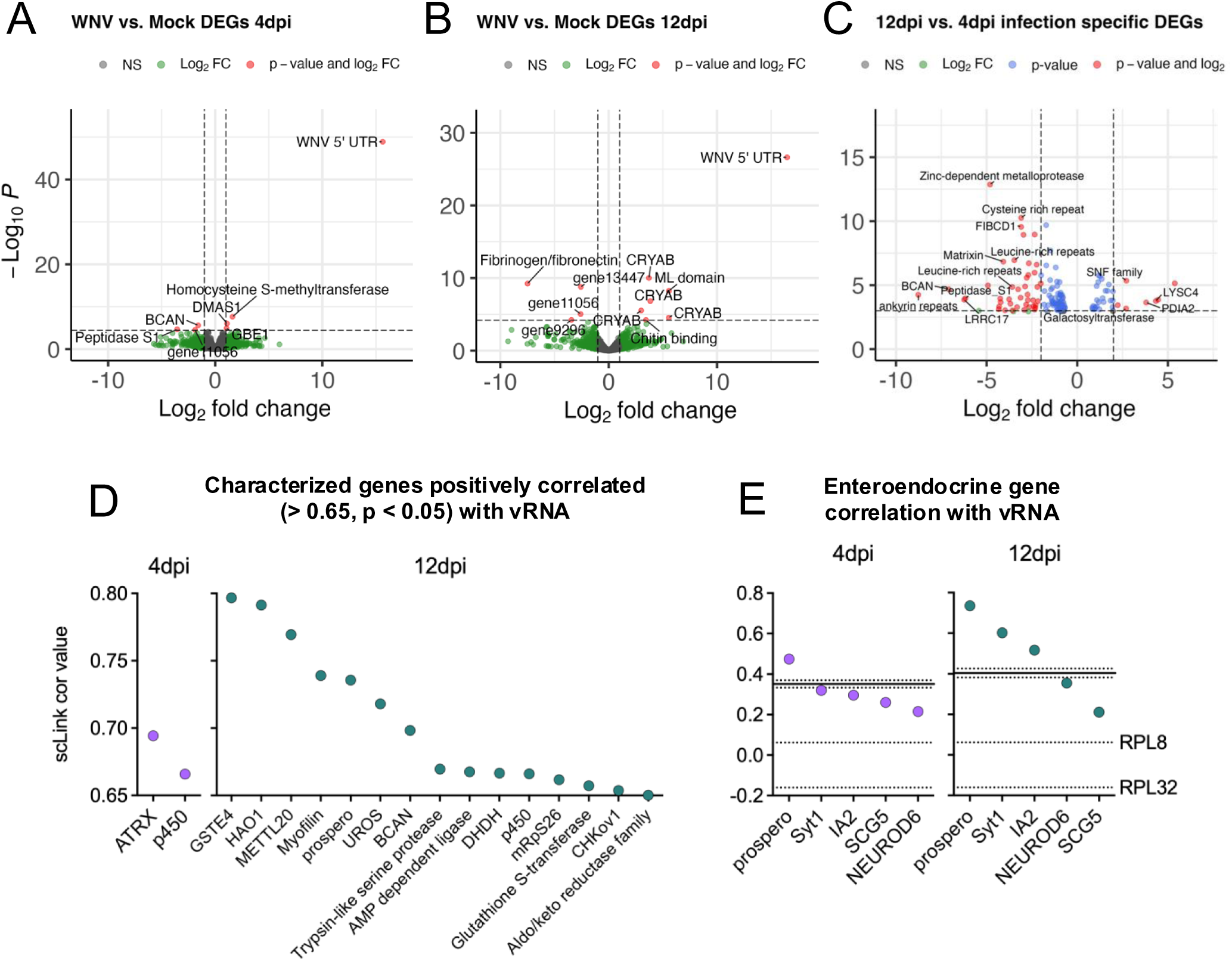
Identifying genes upregulated in response to WNV infection with differential expression and correlation analyses. DEGs associated with WNV-infected condition when compared to the mock condition at 4dpi (**A**) and 12dpi (**B**) were identified with DESeq2. Negative log_2_FC indicates downregulation and positive log_2_FC indicates upregulation associated with WNV infection. (**C**) Genes upregulated during WNV infection at 4dpi (negative log_2_FC) and 12dpi (positive log_2_FC). (**D**) Characterized genes that are significantly correlated with WNV vRNA at 4dpi (purple) and 12dpi (teal), with correlation values of >0.65. Only significant (p < 0.05) relationships shown. (**E**) Correlation of enteroendocrine specific genes and WNV vRNA at 4dpi (purple) and 12dpi (teal). Unlabeled solid line represents the mean correlation of WNV vRNA with 1000 randomly selected genes at the specified timepoint (unlabeled dotted lines represent the upper and lower 95% confidence intervals for this value). Labeled dotted lines denote vRNA correlation with *RPL8* and *RPL32* housekeeping reference genes. Panels C-E were derived from only WNV-infected replicates.

To further examine genes associated with WNV infection, we performed a gene correlation on normalized counts for the top 500 variable genes (genes that have variation in expression across all cells) for each timepoint, determined significance via bootstrapping, then extracted and visualized characterized genes correlated (>0.65) with vRNA (**Figure 5D**, **Supplemental Table 4.2**). Transcription regulator *ATRX*, a cytochrome p450 gene and several uncharacterized genes were strongly correlated with vRNA at 4dpi (**Figure 5D, Supplemental Table 4.2**). At 12dpi, *GSTE4*, *HAO1*, *METTLE20*, *prospero*, *UROS*, *BCAN*, *DHDH*, and *CHKov1* were strongly correlated with vRNA along with serine protease, AMP dependent ligase, cytochrome p450, mitochondrial ribosomal S26, glutathione S-transferase, and aldo/keto reductase family genes (**Figure 5D, Supplemental Table 4.2**). Many of these genes have no documented roles in flavivirus infection.

However, *ATRX* has been implicated in the cellular response to DNA damage, and chromatin remodeling – processes which many viruses exploit during infection (43–45). Further, cytochrome p450 enzymes and serine proteases have been purported to play a role in the mosquito response to viral infections (46–49).

Upon observing that *prospero*, the canonical marker for EE cells, was significantly positively correlated with vRNA at 12dpi, we examined the correlation between vRNA and several previously described neuroendocrine genes (**Figure 2D**, **5E**). Two previously described housekeeping genes, *RPL8* and *RPL32* (50), were validated as having broad expression throughout the total population and used to both confirm that the high prevalence of vRNA in these populations was not confounding the results and provide a visual reference for a biologically insignificant correlation value (**Figure 5E**). Additionally, for each timepoint we determined the correlation between WNV vRNA and 1,000 random genes from the dataset (unlabeled solid line denotes the average of this calculation with 95% confidence intervals) (**Figure 5E**). At 4dpi *prospero*, and at 12dpi *prospero*, *Syt1*, and *IA2* had positive correlations with vRNA (**Figure 5E**).

### Characterization of the midgut immune response to WNV infection at the whole-tissue and single-cell level

While previous work demonstrated an increase in hemocyte proliferation upon bloodmeal ingestion and infection, there were no significant increases in the proportion of hemocyte populations associated with infection at either time point (**Figure 6A-B**) (51,52). Upon observing that no mosquito immune genes were identified as significantly upregulated in response to WNV infection by pseudo-bulk DEG and correlation analyses, we manually compared percent of cells expressing and expression levels of key immune genes that have been implicated in viral control/infection response (19,37,53–56). We identified orthologs in the *Cx. tarsalis* genome for mosquito immune genes; *DOME*, *NANOS1*, *MYD88*, *IMD*, *AGO2*, *R2D2*, *STAT*, *Cactus*, *PIAS1*, *SUMO2*, *LYSC4*, *MARCH8*, *PIWI*, and *REL* (**Supplemental Table 5**) and found no significant differences in the percent of cells expressing or average expression of these genes at either time point (**Figure 6C-D**). Next, we examined the relationships between expression of these immune genes and vRNA at the single-cell level in the WNV-infected population (**Figure 6E**). Interestingly, almost all genes were significantly positively correlated with vRNA at both timepoints (**Figure 6E**). To further confirm these findings, we visualized the relationship of the four most highly correlated immune genes (*IMD*, *PIWI*, *PIAS1*, and *DOME*) with vRNA in individual cells, and compared the expression level of each immune gene in both mock and WNV-infected conditions (timepoints combined) (**Figure 6F-G, Supplemental Figure 10**). These genes and vRNA were correlated, despite comparable expression levels of each gene across infection conditions, confirming that while vRNA load can be correlated with specific immune genes at the individual cell level, it does not induce significant immune gene enrichment at the total population level (**Figure 6F-G, Supplemental Figure 10**).

**Figure 6.**
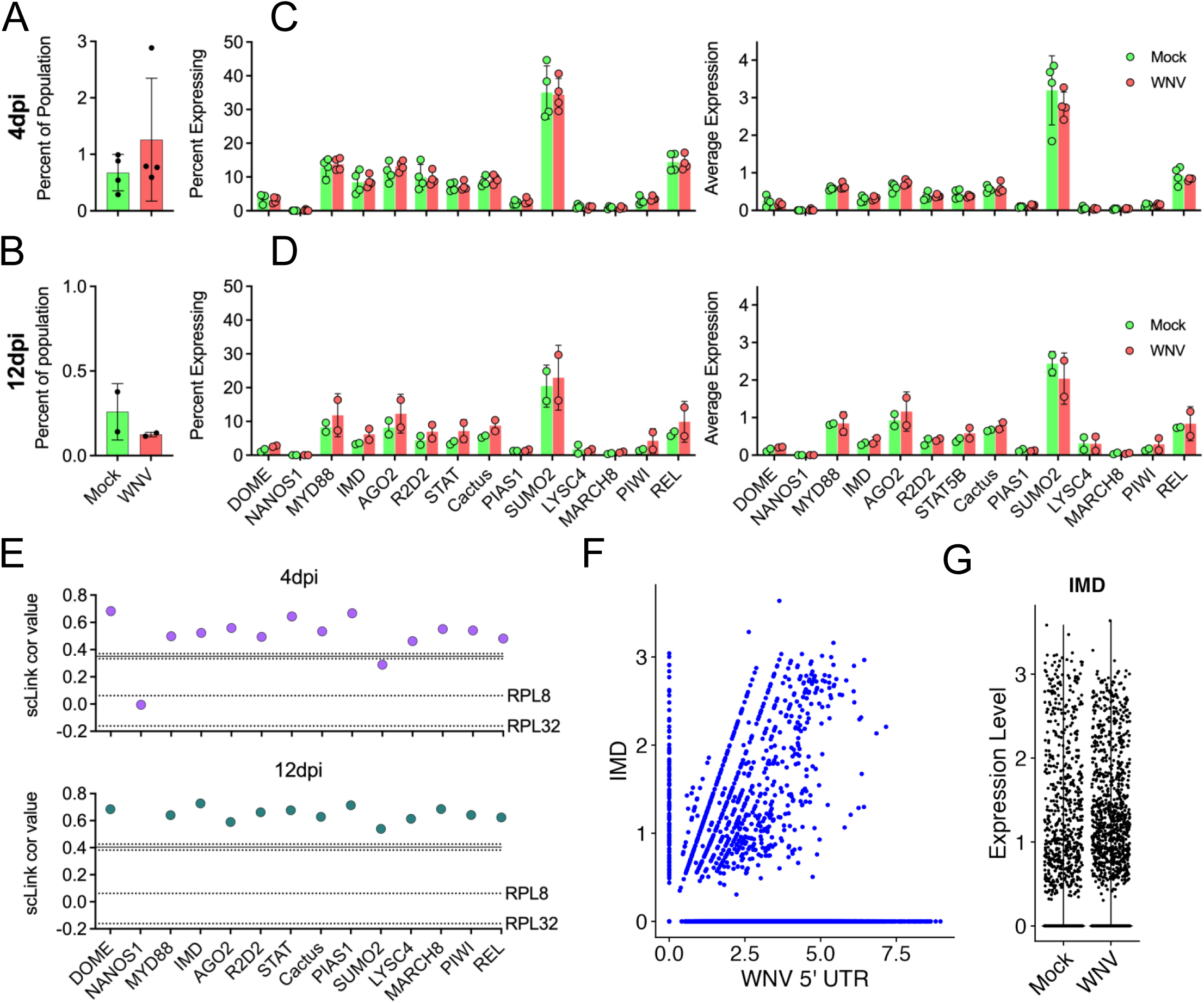
Lack of key immune gene upregulation in response to WNV infection. (**A-B**) Percent of the total population of each replicate comprised by hemocytes compared between mock and WNV-infected conditions at each timepoint. Significance was determined by unpaired t-test. Points = replicate values, bars = mean of replicate values, error bars = SD. (**C-D**) Average expression of, and percent of cells expressing, mosquito innate immune genes compared between mock and WNV-infected replicates for each timepoint. Significance was determined by multiple unpaired t-tests. Points = replicate values, bars = mean of replicate values, error bars = SD. (**E**) Correlation of immune genes and WNV vRNA at 4dpi (purple) and 12dpi (teal). Unlabeled solid line represents the mean correlation of WNV vRNA with 1000 randomly selected genes at the specified timepoint (unlabeled dotted lines represent the upper and lower 95% confidence intervals for this value). Labeled dotted lines denote vRNA correlation with RPL8 and RPL32 housekeeping reference genes. (**F**) Feature scatter of IMD vs. WNV 5’ UTR for both timepoints combined. (**G**) Expression level of IMD in mock and WNV-infected conditions for both timepoints combined. Panels E-F were derived from only WNV-infected replicates.

## Discussion

In this study, we sought to generate a midgut cell atlas (i.e., map of cell type and function) for *Cx. tarsalis* and characterize WNV infection of the midgut at single-cell resolution by performing scRNA-seq on mock and WNV-infected midguts, collected at days 4 and 12 post infection. We identified and described nutrient absorptive (enterocyte), secretory (enteroendocrine), peritrophic matrix secreting (cardia), undifferentiated progenitor (intestinal stem cell/enteroblast), visceral muscle, immune (hemocyte), and fat body cell populations (5,17,18). The distribution and proportion of each cell-type in the total population varied between timepoints, however we identified at least one cluster comprised of each cell-type at each timepoint. Several clusters were precluded from identification due to either lack of canonical markers/enrichment patterns, or origination from a single replicate. Nonetheless, we have demonstrated that single-cell sequencing of *Cx. tarsalis* midguts is feasible and that distinct cell populations can be identified and characterized using previously described canonical cell-type markers and enrichment patterns (14,16–18,20).

There were several differences between our identification of populations in the *Cx. tarsalis* midgut and previously described cell typing efforts in the midguts of *Ae. aegypti* and *Drosophila* (14,17,22,23). Most notably, *Delta*, the canonical marker gene for intestinal stem cells (ISCs) in both *Ae. aegypti* and *Drosophila*, was not enriched in *Cx. tarsalis* ISCs which were, instead, identified by their colocalization with enteroblasts (EBs – enterocyte progenitors) within the UMAP projection (14,17). While *klumpfuss*, the EB marker gene, was specifically expressed in ISC/EB populations, we saw only nonspecific expression of *piezo*, the marker for enteroendocrine progenitors in *Drosophila* (**Supplemental Figure 12**) (14). Further, while there is reference to ISC differentiation markers and proliferative capabilities, existing midgut cell atlases for *Ae. aegypti* and *Drosophila* have not identified a distinct subset of proliferating ISCs – a cell population that, in *Cx. tarsalis*, is highly transcriptionally active and appears important in the context of virus infection (**Supplemental Figure 6**) (14,17,22,23). *Nubbin*, the canonical marker gene for enterocytes (ECs), was reported to be a marker for both EC and EC-like populations in *Ae. aegypti* (17). In our dataset *nubbin* was present in ECs and EC-like cells but was solely identified as a marker in the EC population, while EC-like populations were identified by high expression of a variety of digestive enzyme genes. Additionally, *gambicin* and *pgant4*, previously described cardia cell markers, were not enriched in *Cx. tarsalis* cardia cell populations (14,17,22,23). No *gambicin* ortholog was identified in the *Cx. tarsalis* gene-set despite the existence of a *gambicin* ortholog in the *Cx. quinquefasciatus* genome (CPIJ016084) and, while several *pgant4* orthologs were identified in this gene-set, they were not specifically expressed in any cell population (**data not shown**). Thus, identification of cardia populations in our system was based on enrichment for chitin-binding genes, sugar transport genes, and digestive enzyme genes similar to those found in EC-like populations. Our enteroendocrine (EE) cell population was identified by expression of the canonical marker *prospero*, which aligns with descriptions of EE identification in other systems. However, our EE population showed little to no expression of the previously described mosquito and *Drosophila* neuropeptides/neuropeptide receptors (14,17,20,30,32). This could be due to the known bias of scRNA-seq towards highly expressed genes, or due to *Cx. tarsalis* EE cell secretion of yet uncharacterized neuropeptides. Finally, *NEUROD6* – an ortholog for a neurogenic differentiation factor frequently found in neurons involved in behavioral reinforcement – was highly expressed in our EE population (36). Similarly, a gene commonly expressed in neurons (*CNMa*) has been identified in a subset of EE cells within the *Drosophila* midgut (14).

We detected WNV RNA (vRNA) in the majority of midgut cells at both timepoints. While vRNA significantly increased in the total midgut by 12dpi, the percent of infected cells did not, demonstrating that the majority of midgut cells that will become infected are infected by 4dpi, while vRNA load increases as infection progresses. The high percentage of WNV-infected cells and permissibility of most cell populations to infection supports previous work demonstrating the extreme vector competence of *Cx. tarsalis* for WNV (9,57–59).

Interestingly, while WNV infected almost all midgut cell populations, cluster 4 was significantly enriched for vRNA at both timepoints. This high WNV-expressing cluster was associated with very few (<15) defining cluster markers, precluding us from identifying its cell-type. The cluster markers associated with this cell state are composed entirely of mitochondrial genes and mitochondrial tRNAs, suggesting a heightened state of cell stress and/or energy production. Importantly, this cluster was present and enriched for the same mitochondrial genes in the mock condition, indicating that viral replication did not induce significant stress and/or energy production responses (60,61). Indeed, the presence of cluster 4 in both mock and WNV-infected conditions suggests that this cell state may be associated with blood meal consumption (62,63). Several previous studies have demonstrated that positive sense-single stranded RNA viruses like dengue virus (DENV), and SARS-CoV-2 modulate mitochondrial dynamics to facilitate replication and/or immune evasion (61,64,65), suggesting potential beneficial interactions between WNV and the mitochondria. Additionally, a study in *Lepidoptera* (moths and butterflies) purported that enrichment of mitochondrial genes is associated with insect stress resistance (60) and we demonstrated that several heat shock genes (known to be protective against cell stress) were significantly upregulated in WNV-infected midguts, suggesting possible interplay between WNV infection and the stress response (66–68). However, these heat shock genes were mainly enriched in cluster 8, an un-typed cluster that does not contain the high level of vRNA seen in cluster 4. Additionally, the absence of apoptotic markers in cluster 4 does not discount the possibility that mitochondrial genes are heightened in these cells because they are in the process of dying. If these are dying cells, increased membrane permeability is to be expected and thus high levels of vRNA could be an artifact of the cell state, not a result of it (63).

At 4dpi the EE population is trending toward having more vRNA than other epithelial cell populations and, by 12dpi, has significantly higher levels of vRNA than other populations. Prospero (the canonical marker for EE cells) and select neuroendocrine and vesicle docking genes present in EE cells were positively correlated with vRNA by 12dpi. Additionally, two innate immune system genes were significantly downregulated in the WNV-infected EE population compared to mock at 4dpi. These findings led us to hypothesize that that EE cells serve as sites of enhanced WNV replication during midgut infection. This hypothesis is further supported by previous studies suggest arboviruses preferentially replicate in highly polarized cell types, such as EE cells (69,70), and previous work with Sindbis virus (SINV) - a mosquito-borne alphavirus – that identified EE cells as a site of infection initiation in *Ae. aegypti* (71). Further, a recently published scRNA-seq study of Zika virus infection of the *Ae. aegypti* midgut implicated EE cells as potential key players in infection (23).

In contrast, the levels of vRNA in ISC/EB and ISC/EB-prol populations, relative to the total population, were strikingly and significantly low. We identified the highest number of DEGs associated with WNV infection in ISC/EB and ISC/EB-prol populations, indicating that they have a greater transcriptional response to WNV infection than other midgut cell types. The majority of these DEGs were downregulated, which is consistent with previous scRNA-seq studies in *Aedes aegypti* (23). Upon visualizing the DEGs identified in ISC/EB and ISC/EB-prol populations we observed that predominantly ribosomal and translation related genes were downregulated in infected populations. GO enrichment analysis of all DEGs (cell populations/timepoints combined) that were significantly downregulated in response to infection confirmed that the most highly enriched GO terms denote involvement in ribosomal structure and translation.

Progenitor cells proliferate in response to midgut stress in order to replenish midgut cells impacted by the stressor (72). While the low levels of vRNA found in ISC/EB and ISC/EB-prol populations could be due to the relative “newness” of the cells comprising this population, our findings suggest that the transcriptional state of ISC/EB populations impedes WNV replication. This is further supported by our cluster of orthologous genes (COG) analysis, which revealed that the base transcriptome of the ISC/EB-prol population – which contains less vRNA than the ISC/EB population and the least vRNA of any epithelial cell population – is composed of fewer COGs that denote translation/ribosome structure and biogenesis than non-proliferating populations.

Further, the downregulation of these genes in the WNV-infected condition compared to the mock condition implies that this unfavorable transcriptional state may be a direct response of ISC/EBs to WNV infection. We are not the first to find evidence that proliferating ISCs play a role in infection control in the midgut. Previous work examining *Plasmodium* infection in the *Anopheles* midgut revealed that decreased oocyst survival was associated with enhanced proliferation of progenitor cells (73). Another study characterizing ISC dynamics in response to DENV in the *Ae. aegypti* midgut found that ISC proliferation increased refractoriness to infection, suggesting that cell turnover is an important part of the midgut immune response, and further implicating ISC/EB populations in infection control within the midgut (74).

While the presence of vRNA alone does not signify active replication, average vRNA levels increased between timepoints in all epithelial cell populations, apart from the ISC/EB and ISC/EB-prol populations (**Supplemental Figure 5**), suggesting that the high vRNA levels in EEs, and low vRNA levels in ISC/EBs, are the result of enhanced and restricted replication respectively. The lack of significant differences associated with the rate of ISC/EB and EE infection at both timepoints further supports that varying levels of vRNA in EE and ISC/EB-prol cells is due to permissivity to replication and not susceptibility to infection. The factors that determine cell-type specific enhancement or suppression of WNV replication are not currently known, but could include more efficient evasion of antiviral pathways, more efficient mechanisms of midgut escape/dissemination, abundance of pro/anti-viral genes or, in the case of ISC/EBs, scarcity of specific genes related ribosome structure and translation.

We captured a population of sessile hemocytes (hemocytes attached to tissue) that we identified as mature granulocytes – a class of hemocytes known to play an important role in the phagocytic and lytic components of the mosquito immune response – at both timepoints, allowing us to characterize a cellular immune component of mosquito midguts (5,15). The classification of this population as granulocytes was unsurprising, given estimates that granulocytes comprise more than 85% of hemocyte populations in the adult mosquito (51).

There was no evidence of increased immune cell proliferation and attachment to the midgut, or immune gene upregulation in the total infected population compared to mock – suggesting little to no immune activation in the midgut upon WNV infection. This was surprising given the importance of the midgut as a site of innate immune activation (5,6,58,59). Although scRNA-seq of *Cx. tarsalis* after WNV infection has not been described, several previous studies in *Ae. aegypti* and *Cx. pipiens* have noted significant upregulation of IMD and Toll pathway genes in response to viral infection and highlighted that the innate immune response in mosquitoes is a strong determinant of vector competence (66,75,76). The absence of a notable immune response to WNV infection in *Cx. tarsalis* could be a determinant of the vector’s extreme susceptibility and competence (9,57–59). However, in individual cells, most immune genes had some degree of significant positive correlation with vRNA suggesting that, while WNV infection does not cause significant enrichment of these genes in the total population, WNV infection and replication can influence the expression of these genes at the single-cell level. This finding highlights that scRNA-seq is a powerful tool for characterizing infection dynamics that are not apparent when looking at the population average.

### Limitations and Future Directions

Our inability to detect certain genes (i.e., neuropeptide genes and additional canonical markers we would expect to see) could be due to the low percentage (∼30%) of reads mapped to the *Cx. tarsalis* genome (likely due to either the presence of reads that were too short to map, or the current state of the *Cx. tarsalis* genome annotation), or the absence of those genes in the existing annotation file. Future scRNA-seq studies in *Culex* mosquitoes could potentially benefit from adjusting the fragmentation time recommended by 10X Genomics. Further, improvement of the existing *Cx. tarsalis* genome annotation would facilitate studies of gene expression in this species. A multitude of genes detected in our dataset remain uncharacterized due to a lack of appropriate orthologs, which could be explained by the evolutionary divergence between *Cx. tarsalis* and the species from which most gene orthologs were derived; *Cx. quinquefasciatus* and *Ae. Aegypti,* which diverged 15-22 million years ago (MYA) and 148-216 MYA respectively (12). High levels of vRNA in specific cell types imply that replication is occurring/has occurred but it is important to note that the presence of vRNA is not analogous to active viral replication (e.g., the presence of vRNA could be the result of phagocytosis of an infected cell). Future studies could use qRT-PCR, focusing on our top genes of interest, to measure expression kinetics and levels following midgut infection in *Cx. tarsalis* and other relevant vectors. Finally, early WNV infection in the *Cx. tarsalis* midgut will be further studied in our lab via immunofluorescence and flow cytometry assays using cell-type and cell state specific antibodies in whole midguts, putting the findings described here into a spatial context.

## Conclusion

The work presented here demonstrates that WNV is capable of infecting most midgut cell types in *Cx. tarsalis*. Moreover, while most cells within the midgut are susceptible to WNV infection, we observed modest differences in virus replication efficiency that appeared to occur in a cell-type specific manner, with EE cells being the most permissive and proliferating ISC/EB cells being the most refractory. Further, our findings demonstrate that ISC/EBs downregulate genes associated with ribosome structure and translation during WNV infection, and suggest that the resulting transcriptional state of ISC/EBs is unfavorable for WNV replication. We observed mild to no upregulation of key mosquito immune genes in the midgut as a whole, however, we show that immune gene expression can be correlated with WNV vRNA level within individual cells. Additionally, we have generated a midgut cell atlas for *Cx. tarsalis* and, in doing so, improved the field’s understanding of how WNV establishes infection in this highly competent vector.

## Acknowledgements

The research reported in this publication was supported by National Institutes of Health grant R01-AI067380 (GDE), National Institute of Allergy and Infectious Diseases of the National Institutes of Health under Award Number T32-AI162691 (EAF), and by Colorado State University’s Office of the Vice President for Research’s “Accelerating Innovations in Pandemic Disease” initiative, made possible through support from The Anschutz Foundation (EAF). Emily N Gallichotte was supported by funding to Verena (viralemergence.org) from the U.S. National Science Foundation, including NSF BII 2021909 and NSF BII 2213854. The content is solely the responsibility of the authors. Next Generation Sequencing was performed at the Genomics Shared Resource (RRID: <u>SCR_021984</u>) at the University of Colorado. This resource is supported by Cancer Center Support Grant (P30CA046934). MKNL, SP, and VH were all supported by the Wellcome Trust (grant 206194/Z/17/Z) and MKNL was supported by an MRC Career Development Award G1100339. The funders had no role in study design, data collection and analysis, decision to publish, or preparation of the manuscript.

## Author Contributions

Emily A. Fitzmeyer – Conceptualization, validation, formal analysis, investigation, resources, data curation, writing – original draft, writing – review and editing, visualization, project administration, funding acquisition. Taru S. Dutt – Validation, resources, investigation, supervision, writing – review and editing.

Silvain Pinaud – Methodology.

Barb Graham – Formal analysis.

Emily N. Gallichotte – Visualization, investigation, writing – review and editing.

Jessica Hill – Validation, resources.

Corey Campbell – Validation.

Hunter Ogg – Resources, investigation.

Virginia Howick – Methodology.

Mara Lawniczak – Supervision.

Erin Osborne Nishimura – Validation, supervision.

Sarah Helene Merkling – Validation, resources.

Marcela Henao-Tamayo – Validation, resources, supervision.

Gregory D. Ebel – Conceptualization, validation, resources, writing – review and editing, supervision, funding acquisition.

## Inclusion and diversity

We support inclusive, diverse, and equitable conduct of research.

## Methods

### Virus

All infections were performed with a recombinant barcoded WNV (bcWNV) passage 2 stock (epidemic lineage I strain, 3356) grown on Vero cells. Titer for the stock was determined by standard Vero cell plaque assay (77).

### Mosquito infection

Mosquito studies were conducted using laboratory colony-derived *Cx. tarsalis* mosquitoes (>50 passages). WNV infections in mosquitoes were performed under A-BSL3 conditions. Larvae were raised on a diet of powdered fish food. Mosquitoes were maintained at 26°C with a 16:8 light:dark cycle and maintained at 70–80% relative humidity, with water and sucrose provided ad libitum. *Cx. tarsalis* mosquitoes were transferred to A-BSL3 conditions 48 hours prior to blood feeding, and dry starved 20-24 hours before blood feeding. Seven days after pupation (6-7 days after emergence) mosquitoes were exposed to an infectious bloodmeal containing a 1:1 dilution of defibrinated calf’s blood and bcWNV stock diluted in infection media (Dulbecco’s Modified Eagle’s Medium, 5% penicillin-streptomycin, 2% amphotericin B, and 1% fetal bovine serum (FBS)) for a final concentration of 3-6e^7^ PFU/mL, or a mock bloodmeal containing a 1:1 dilution of defibrinated calf’s blood and infection media. All bloodmeals were provided in a hog’s gut glass membrane feeder, warmed by circulating 37°C water. Following 50-60 minutes of feeding, mosquitoes were cold-anesthetized, and engorged females were separated into cartons and maintained on sucrose.

### Collection of mosquito tissues

At indicated time points, mosquitoes were cold-anesthetized and transferred to a dish containing Sf900III insect cell culture media (Gibco) with 5% FBS. Midguts were dissected, and transferred to tubes containing 500uL Sf900III media + 5% FBS and kept on ice for the duration of dissections. Ten pooled midguts per sample were collected for dissociation and sequencing.

### Midgut dissociation and single-cell suspension preparationi

A dissociation buffer containing *Bacillus licheniformis* protease (10mg/mL) and DNAse I (25U/mL) was prepared in Sf900III media (Gibco). Pools of 10 midguts were resuspended in dissociation media, transferred to a 96-well culture dish, and triturated with a p1000 pipet at 15-20 minute intervals for 105 minutes. At each interval 100-125μ L containing dissociated single-cells was collected (with replacement) from the top of the dissociation reaction and transferred to 25mL of Sf900III + 5% FBS on ice. Dissociation reactions were kept covered at 4°C between trituration. Upon complete tissue dissociation the entire remaining volume of each reaction was transferred to Sf900III + 5% FBS on ice. Collection tubes with dissociated cells were centrifuged at 700xg for 10 minutes at 4°C, resuspended in 500μL of Sf900III + 5% FBS, and passed through a 40μ small volume filter (PluriSelect). Immediately prior to loading on the Chromium Controller, cell suspensions were spun down at 700xg for 10 minutes and washed twice in 1mL PBS + 0.04% bovine serum albumin (BSA), and resuspended in 50μL of PBS + 0.04% BSA. Cell concentration and viability was determined using the Countess II Automated Cell Counter (Thermo Fisher Scientific). Appropriate cell suspension volume (target recovery of 10,000 live cells) was loaded on the Chromium controller. Further details on this protocol can be found here - dx.doi.org/10.17504/protocols.io.j8nlke246l5r/v1.

### Gel Bead-In Emulsions (GEM) generation and cDNA synthesis

GEM generation and cDNA synthesis were performed using the Next GEM Single Cell 5’ GEM kit v2 (PN-1000266) and Next GEM Chip K Single Cell Kit (PN-1000286) (10X Genomics). Reactions for GEM generation were prepared according to the Chromium Next GEM Single Cell 5’ Reagent Kit v2 (dual index) user guide with one alteration; a primer specific to the WNV envelope region of the genome (5′-AAGCAGTGGTATCAACGCAGAGTAC-GGGTCAGCACGTTTGTCATTG-3′) was added to all samples at a concentration of 10nM (7.5μl – displacing l of the total H_2_O added to each reaction) (21,78). GEMs were generated using the 10X Chromium controller X series and cDNA was synthesized according to the Chromium Next GEM Single Cell 5’ Reagent Kit v2 (dual index) user guide.

### Library preparation and sequencing

Four libraries were prepared using the Next GEM single cell 5’ v2 library construction kit and Dual Index Kit TT Set A (10X Genomics, PN-1000190 and PN-1000215 respectively). Library construction was carried out according to the Chromium Next GEM Single Cell 5’ Reagent Kit v2 (dual index) user guide. Library concentration was determined by KAPA Library Quantification Kit (Roche). Libraries were then diluted to 15nM, pooled by volume, and sequenced at the CU Anschutz Genomics and Microarray core on the NovaSeq 6000 (150x10x10x150) (Illumina) at a target coverage of 4.0e^8^ read pairs per sample, equating to 40,000 read pairs per cell. Average sample coverage and cell recovery per sample was 5.3e^8^ read pairs and 2417.9 cells respectively (**Supplemental File 1**).

### Reference generation and sample processing with Cell Ranger

The gene feature files associated with the *Cx. tarsalis* and WNV genomes were converted to gene transfer format (gtf) using AGAT (v1.0.0) prior to being filtered with the CellRanger (v7.0.1) mkgtf function (79). An ‘MT-‘ prefix was manually added to all non-tRNA features located in the mitochondrial chromosome of the *Cx. tarsalis* genome. The contents of the filtered WNV genome feature file, and fasta file, were appended to the *Cx. tarsalis* feature and fasta files respectively, and run through CellRanger::mkref. All sequencing data were processed and mapped to the aforementioned *Cx. tarsalis*/WNV combined reference genome using CellRanger::count with the following parameters: --include-introns=true \ --expect-cells=10000.

### Quality control and Seurat workflow

Cell Ranger output files were individually read into RStudio (RStudio - v2023.09.0+463, R – v4.3.2) as SingleCellExperiment objects using the singleCellTK package (v2.12.0).

Doublet identification and ambient RNA estimation were performed with singleCellTK::runCellQC using the algorithms “scDblFinder” and “DecontX” respectively. Samples were filtered for doublets and ambient RNA contamination by keeping cells with the following metrics: decontX_contamination < 0.6, scDblFinder_doublet_score < 0.9. Samples were then converted to Seurat objects, log normalized individually, and merged into one Seurat object, with columns pertaining to sample of origin, infection condition, and timepoint added to the object metadata prior to processing as described in the Seurat (v4.3.0.1) guided clustering tutorial (80). Briefly, mitochondrial gene percentage for each cell was calculated, and cells with the following metrics were retained: nFeature_RNA > 100, nFeature_RNA < 2500, percent_mt < 25 (14). Cell retention metrics were informed by a previous study of the Drosophila midgut (14). Percent of cells retained after QC for each sample can be found in **Supplemental File 1**. Features were log normalized, variable features were identified, the top 2000 variable features were scaled, a principal component (PC) analysis dimensionality reduction was run, and the number of PCs needed to adequately capture variation in the data was determined via elbow plot. Nearest neighbors were computed, and appropriate clustering granularity was determined with Clustree (v0.5.0). Uniform Manifold Approximation and Projection (UMAP) dimensional reduction was performed, and clusters were visualized with UMAP reduction. Cluster markers were identified with Seurat::FindConservedMarkers using default parameters and infection condition as the grouping variable. WNV vRNA as a cluster marker was identified by splitting the dataset by infection condition and timepoint and using Seurat::FindAllMarkers on the WNV-infected samples with a logfc threshold of 0.25. In all cases where calculations were performed on individual replicates or individual conditions, the merged Seurat object was split by sample or condition using Seurat::SplitObject so that calculations performed on subsets of the data or individual replicates were derived from a dataset that had been normalized as one. Percent expression and average expression were calculated with scCustomize::Percent_Expressing (v1.1.3) and Seurat::AverageExpression respectively. Feature expression levels were visualized with Seurat::FeaturePlot and Seurat::VlnPlot. Solely mitochondrial genes and tRNAs were identified as markers for cluster 4, suggesting that this cluster was composed of dying cells. While it is common practice to remove dying cells from scRNAseq datasets further examination of cluster 4 revealed 12.27% of features were mitochondrial, 0.97% were viral, and 33.91% were tRNA, demonstrating that the percentage of mitochondrial genes was well below our QC threshold and was not artificially deflated by high vRNA levels. As we could neither confirm cell death nor rule out the possibility that cluster 4 represents a highly metabolically active cell state we did not see fit to remove this population from our dataset.

### Trajectory inference with Slingshot

Lineage structure and pseudotime inference was performed using the Slingshot (v2.10.0) and tradeSeq (v1.16.0) functions getLineages, getCurves, and fitGAM successively on the dataset containing epithelial cell populations (timepoints combined). ISC/EB and ISC/EB-prol populations were specified as the start state and EC and EE populations were specified as the end state for lineage determination.

### Pseudo-bulk differential expression analysis with DESeq2

Pseudo-bulk dataset was generated and differential expression analysis performed as described by Khushbu Patel’s (aka bioinformagician) pseudo-bulk analysis for single-cell RNA-Seq data workflow tutorial (81). Briefly, raw counts were aggregated at the sample level using Seurat::AggregateExpression, and the aggregated counts matrix was extracted and used to create a DESeq2 object (dds). The dds object was filtered to retain genes with counts >=10 prior to running DESeq2 and extracting results for the appropriate contrast. For total population DEGs specific to timepoints we subset the dataset by timepoint. For DEGs specific to midgut cell populations we subset the dataset by ident/cluster/population prior to performing the DESeq2 workflow. We identified infection associated DEGs between timepoints by performing a pseudo-bulk DE analysis between 4dpi and 12dpi WNV-infected samples and mock samples separately. We then filtered out DEGs associated only with bloodmeal consumption (genes that came up as significantly differentially expressed between timepoints in our mock condition) leaving only DEGs associated with infection. DESeq2 results for the total population were visualized via EnhancedVolcano (v1.20.0). DESeq2 results for specific midgut cell populations were visualized in Prism (10.0.3).

### Gene Ontology Enrichment Analysis

GOEA was performed by generating a list of *Culex quinquefasciatus* ortholog accession numbers corresponding to all up or downregulated DEGs identified in specific cell populations. In instances where accession numbers were available for orthologs in multiple species the *Culex quinquefasciatus* ortholog was selected. Accession number lists were submitted to ShinyGO 0.80 (web version - http://bioinformatics.sdstate.edu/go/) with the *Cx. quinquefasciatus* CpipJ2 assembly set as a reference, FDR cutoff set to 0.05, and default settings for all other parameters (82). GOEA plots were generated using the “chart” tab, and downloaded directly from ShinyGO 0.80 (82).

### Gene correlation with vRNA level

Raw gene counts for each timepoint were extracted and normalized using scLink::sclink_norm (v1.0.1) (83). Correlation matrices were then generated for the top 500 variable genes (identified during the Seurat guided clustering workflow) using scLink::sclink_corr (83). For specific neuroendocrine and immune gene correlations a vector of specific gene names was supplied in lieu of the top 500 variable genes. P-values associated with correlation values were determined via bootstrapping. Average correlation of 1000 random genes was determined by generating a vector of all Seurat object rownames containing the string “gene”, randomly sampling 999 genes from this vector, appending the WNV 5’ UTR feature ID to the subsample, and normalizing and determining correlations as described above. Mean correlation value and 95% confidence intervals were calculated in Prism (10.0.3)

### Ortholog identification with eggNOG mapper

Coding sequence (CDS) genome coordinates were extracted from the *Cx. tarsalis* gene transfer format file and corresponding genome sequences were extracted from the corresponding fasta file using Bedtools (v2.26.0). CDS were then assigned orthologs via eggNOG-mapper v2 (web version - http://eggnog-mapper.embl.de) using default parameters (**Supplemental File 4**) (84,85).

Feature file (gtf) gene IDs were merged with the eggNOG output file using custom R scripts. Gene ID, ortholog seed, preferred gene name, COG category notation, PFAM information, and gene description were retained in a gene name and ID file within which information pertaining to identical gene IDs was aggregated using a custom R script. This aggregated gene name and ID file (**Supplemental File 5**) was used to assign gene names to the marker files used for cell-typing, DESeq2 results files, scLink correlation matrices, etc.

### Statistical analyses

Statistical analyses were performed in GraphPad Prism version 10.0.3. Differences in vRNA level in known clusters were measured by one-way ANOVA with Tukey’s multiple comparisons test. Differences in average expression and percent expression of mosquito immune genes between mock and WNV-infected conditions were measured by multiple unpaired t-tests. Differences in hemocyte population proportion were measured by unpaired t-test.

All scripts used for data processing, analysis, and visualization are available on GitHub: https://github.com/fitz-meyer/scRNA_seq_fitz

RDS files for all samples included in the described analyses are available on GitHub : https://github.com/fitz-meyer/scRNA_seq_rdsFiles.git

All sequencing data and raw matrix files are available upon request.

## Supplemental Material

**Supplemental Figure 1. COG profiles of midgut cell populations.** Cluster of orthologous gene (COG) profiles for (**A**) genes expressed in ≥75% of cells in each cluster and (**B**) cluster marker genes were visualized as percentage of total for each cluster/population. Colors represent COG notation A-Z, NA and DIV as shown in the notation key embedded in the figure. Where applicable, marker gene COG profiles were derived from cluster markers that are conserved between infection conditions.

**Supplemental Figure 2. Cluster proportion and grouping by condition and replicate.** Proportion of the total population comprised by each cluster compared between mock and WNV-infected conditions at 4dpi (**A**) and 12dpi (**B**). Only significant comparisons shown. Significance determined by multiple unpaired t-tests. Bar = mean, error bars = SD. (**C**) Cluster grouping and composition by sample – mock and infected samples both plotted. Different colors denote different samples. (**D**) Cluster grouping and composition by infection condition. Salmon = mock, blue = WNV-infected.

**Supplemental Figure 3. Characterizing EE, ISC/EB, and HC cell-types.** Visualizing expression of neuroendocrine and secretory genes, HC class marker genes, and proliferation and mitotic marker genes across the total population to determine specificity of expression. Violin plots show expression level in individual cells grouped by cluster.

**Supplemental Figure 4.** Enrichment for identical sets of mitochondrial genes and tRNAs specific to cluster 4. Dot size denotes percent of cells expressing each gene, color denotes scaled gene expression. “MT-“ prefix was added to tRNA gene names in this figure for clarity. Mitochondrial tRNAs were not included in mitochondrial gene estimation for QC filtering. Plots were derived from only WNV-infected samples.

**Supplemental Figure 5. Side by side comparison of vRNA levels in midgut cell populations at each timepoint.** Purple = 4dpi, green = 12dpi. Points within bars denote replicate values, bar denotes mean of replicate values. Error bars = SD.

**Supplemental Figure 6. Examining transcriptional activity of clusters.** Feature counts and RNA counts for individual cells within clusters.

**Supplemental Figure 7. Confirming that cell death does not drive clustering.** Expression of apoptotic and anti-apoptotic genes visualized in the total population via UMAP feature plot. Color in feature map denotes expression level.

**Supplemental Figure 8. S phase markers.** Expression of S phase gene markers visualized in the total population via UMAP feature plot. Color in feature map denotes expression level.

**Supplemental Figure 9. G2/M phase markers.** Expression of G2/M phase gene markers visualized in the total population via UMAP feature plot. Color in feature map denotes expression level.

**Supplemental Figure 10. Visually confirming vRNA level is correlated with select immune gene expression without significantly increasing expression in the total population.** Correlation between vRNA and 3 of the most highly correlated immune genes (as determined by scLink) confirmed by feature scatter. Equivalent expression levels of immune genes between mock and WNV-infected conditions confirmed by violin plot. Scatter plots derived from only WNV-infected replicates.

**Supplemental Figure 11. Visualizing cell-type marker expression along epithelial cell lineages identified by trajectory analysis.** Lineage 1 (green – EC lineage) and 2 (blue – EE lineage) displayed expression of klumpfuss (Klu), the canonical marker for enteroblasts, prior to differentiation. PCNA expression was associated with the progression of each lineage into the proliferating ISC/EB population. The canonical marker for ECs (POU2F1/nubbin) was enriched upon differentiation of lineage 1 into ECs. The canonical marker for EEs (PROX1/prospero) was enriched upon differentiation of lineage 2 into EEs.

**Supplemental Figure 12. Nonspecific expression of *piezo*.** Expression of the EEP gene *piezo* in individual cells grouped by cluster.

**Supplemental Table 1. Cluster proportion.** Percent of the total midgut cell population each distinct cluster composes.

**Supplemental Table 2. WNV vRNA as a cluster marker.** Log_2_FC values of the WNV 5’ UTR (nbis-gene-2-utr) in clusters where vRNA was identified as a significant (p-adj < 0.0.5) cluster marker in the WNV-infected condition. Log_2_FC values denote vRNA enrichment in a cluster relative to the total remaining population.

**Supplemental Table 3. All significant DEGs identified as associated with WNV-infection in midgut cell populations.** Significant (p-adj < 0.05) differentially expressed genes (DEGs) grouped by population and timepoint. Log_2_FC values denote enrichment of a gene in a specific cell population in the WNV-infected condition relative to that specific cell population in the mock condition.

**Supplemental Table 4.1 Log_2_FC values and accession numbers for total population pseudo-bulk differential expression analysis.** Gene ID, gene name, Log_2_FC value, ortholog accession number, adjusted p-value, and description for all genes significantly (p-adj < 0.05) differentially expressed between infected and mock conditions.

**Supplemental Table 4.2 Complete list of genes identified as significantly positively correlated with WNV vRNA.** Gene name, ortholog accession number, gene ID, correlation value, p-value, and description for all genes significantly (p < 0.05) positively correlated (> 0.65) with vRNA. Significance was determined by bootstrapping.

**Supplemental Table 5. Innate immune gene identifiers.** Gene ID, gene name, and accession number for mosquito innate immune genes compared between mock and WNV-infected conditions.

**Supplemental File 1. scRNA-seq run summaries.** Provides technical details for all samples processed as part of this study: cells recovered, median genes per cell, mean reads per cell, percent of reads mapped to genome, percent of cells retained after initial QC, percent of cells infected, coverage, %Q30, pooling method, cell viability, prep details/alterations, and justification for exclusion of samples.

**Supplemental File 2. Midgut cell markers.** Gene ID, accession number, name/description, and expression information for all genes used in identifying or supporting the identification of each cell population.

**Supplemental File 3. Top 50 conserved markers.** The top 50 (arranged by positive log_2_FC in the mock condition) conserved cluster markers identified via Seurat’s FindConservedMarkers function for all 20 clusters.

**Supplemental File 4. Original eggNOG mapper output file.** Unaltered eggNOG mapper ortholog identification results for all coding sequences identified in the *Cx. tarsalis* genome feature file.

**Supplemental File 5. Aggregated gene ID ortholog list with accession number/ortholog seed.** eggNOG mapper ortholog identification output merged with gene IDs from the *Cx. tarsalis* genome feature file with gene ID repeats collapsed so that each gene ID corresponds to one ortholog/gene name entry.

