## Supplemental Figures for "A single-cell atlas of the *Culex tarsalis* midgut during West Nile virus infection"

### Slide 1
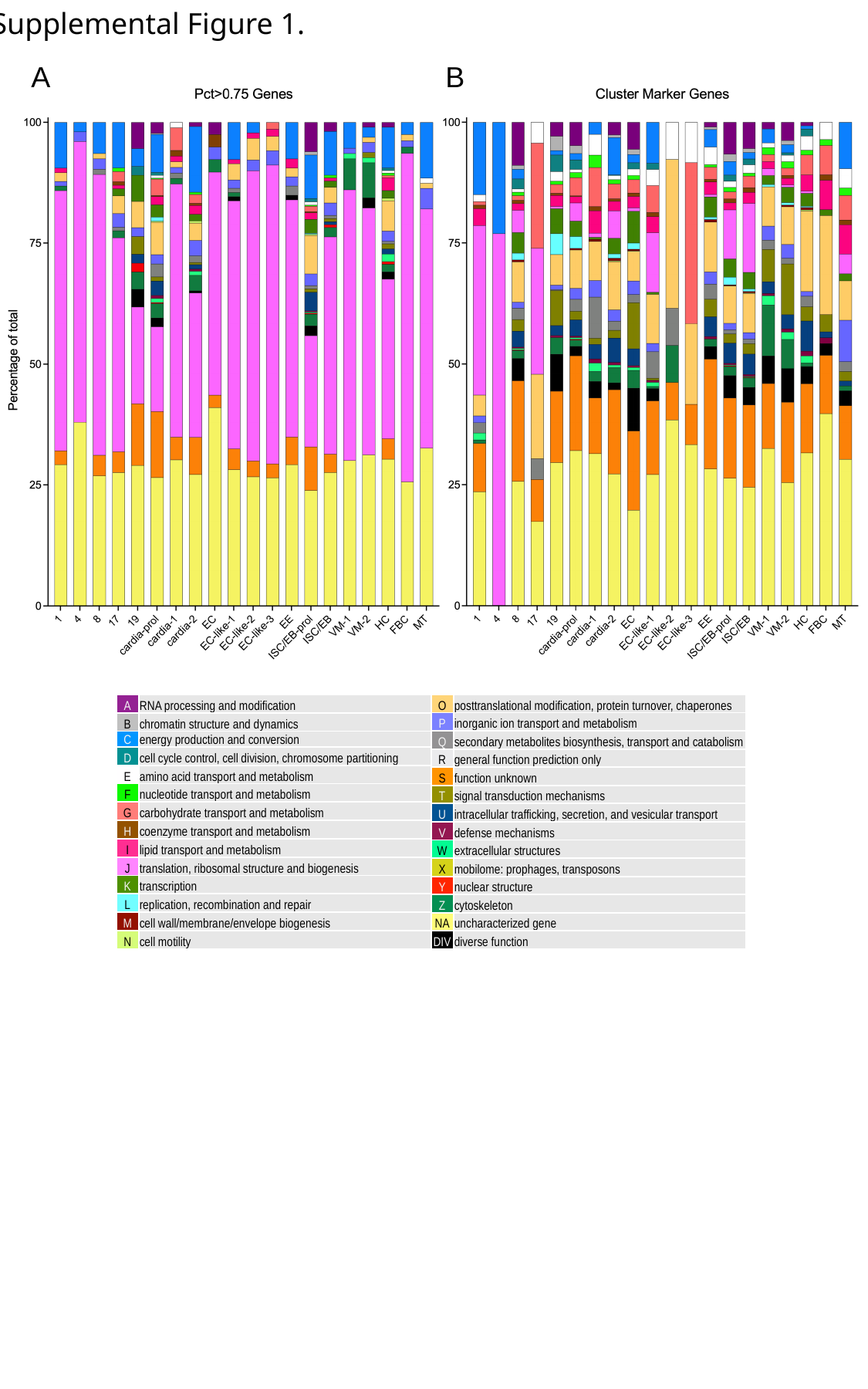

Supplemental Figure 1.
A
B
| A | RNA processing and modification |
| --- | --- |
| B | chromatin structure and dynamics |
| C | energy production and conversion |
| D | cell cycle control, cell division, chromosome partitioning |
| E | ﻿amino acid transport and metabolism |
| F | ﻿nucleotide transport and metabolism |
| G | carbohydrate transport and metabolism |
| H | coenzyme transport and metabolism |
| I | lipid transport and metabolism |
| J | translation, ribosomal structure and biogenesis |
| K | ﻿transcription |
| L | ﻿replication, recombination and repair |
| M | ﻿cell wall/membrane/envelope biogenesis |
| N | ﻿cell motility |
| O | posttranslational modification, protein turnover, chaperones |
| --- | --- |
| P | ﻿inorganic ion transport and metabolism |
| Q | ﻿secondary metabolites biosynthesis, transport and catabolism |
| R | ﻿general function prediction only |
| S | ﻿function unknown |
| T | signal transduction mechanisms |
| U | intracellular trafficking, secretion, and vesicular transport |
| V | defense mechanisms |
| W | extracellular structures |
| X | mobilome: prophages, transposons |
| Y | nuclear structure |
| Z | cytoskeleton |
| NA | uncharacterized gene |
| DIV | diverse function |

### Slide 2
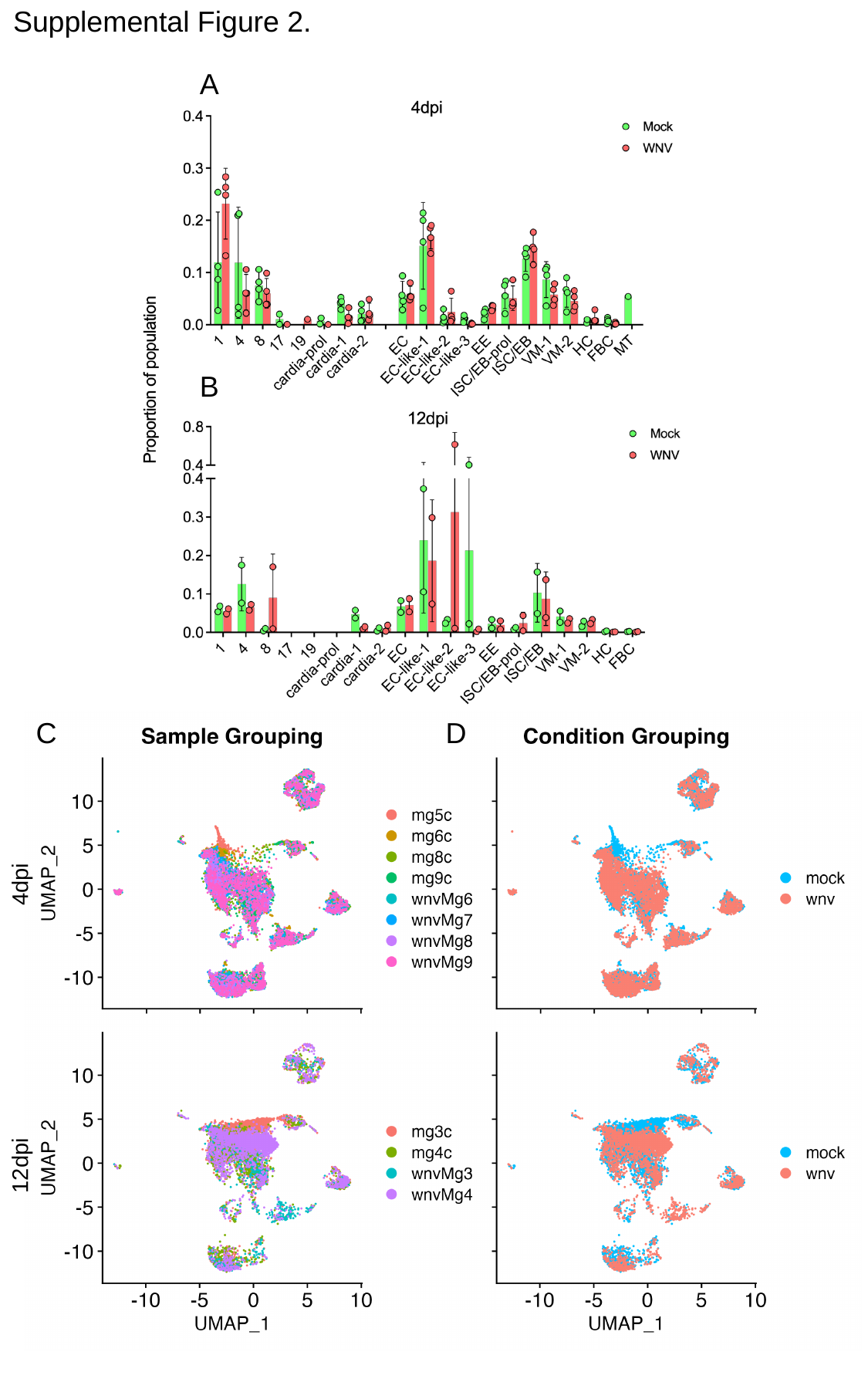

Supplemental Figure 2.
A
B
C
D
4dpi
12dpi

### Slide 3
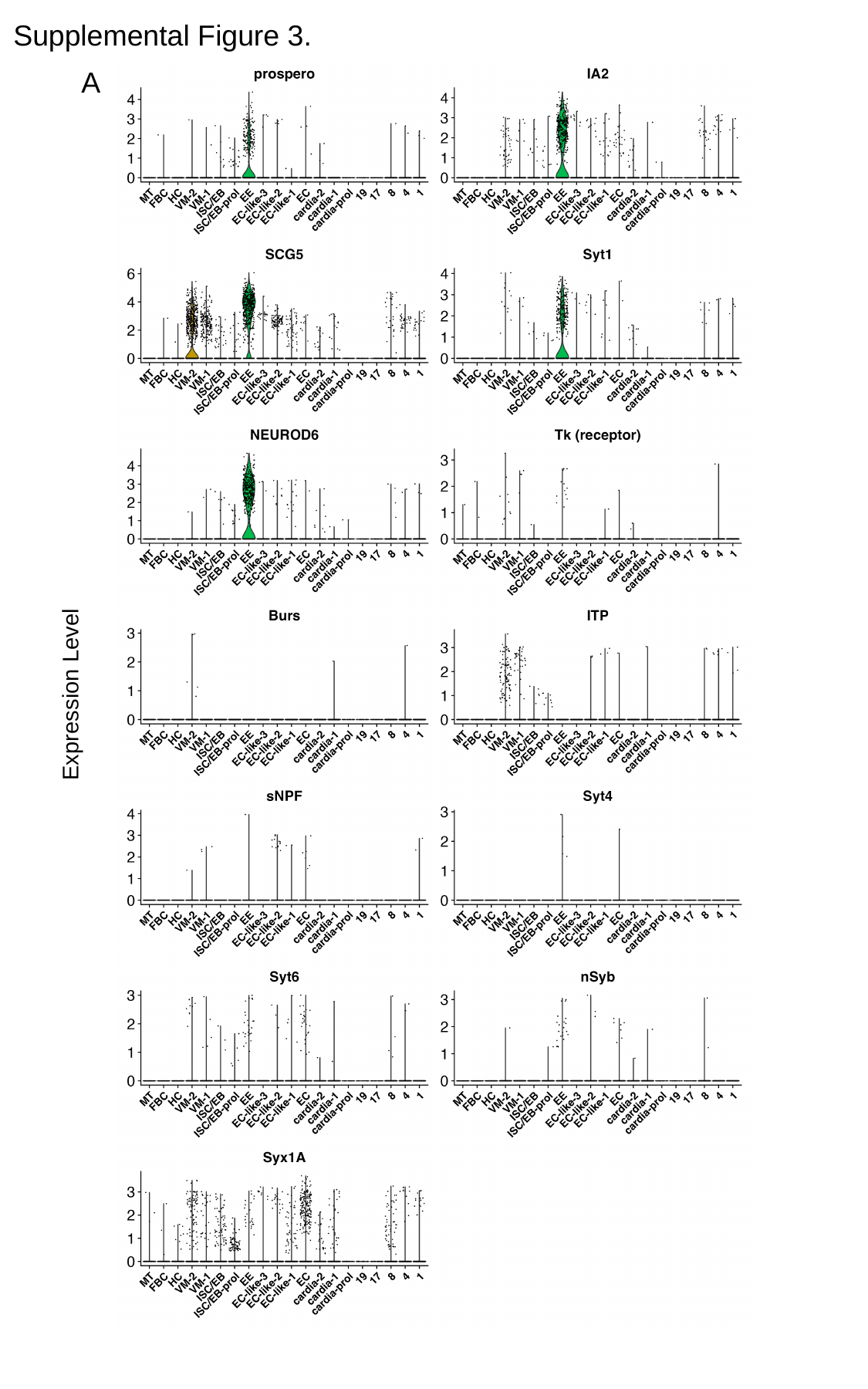

Supplemental Figure 3.
A
Expression Level

### Slide 4
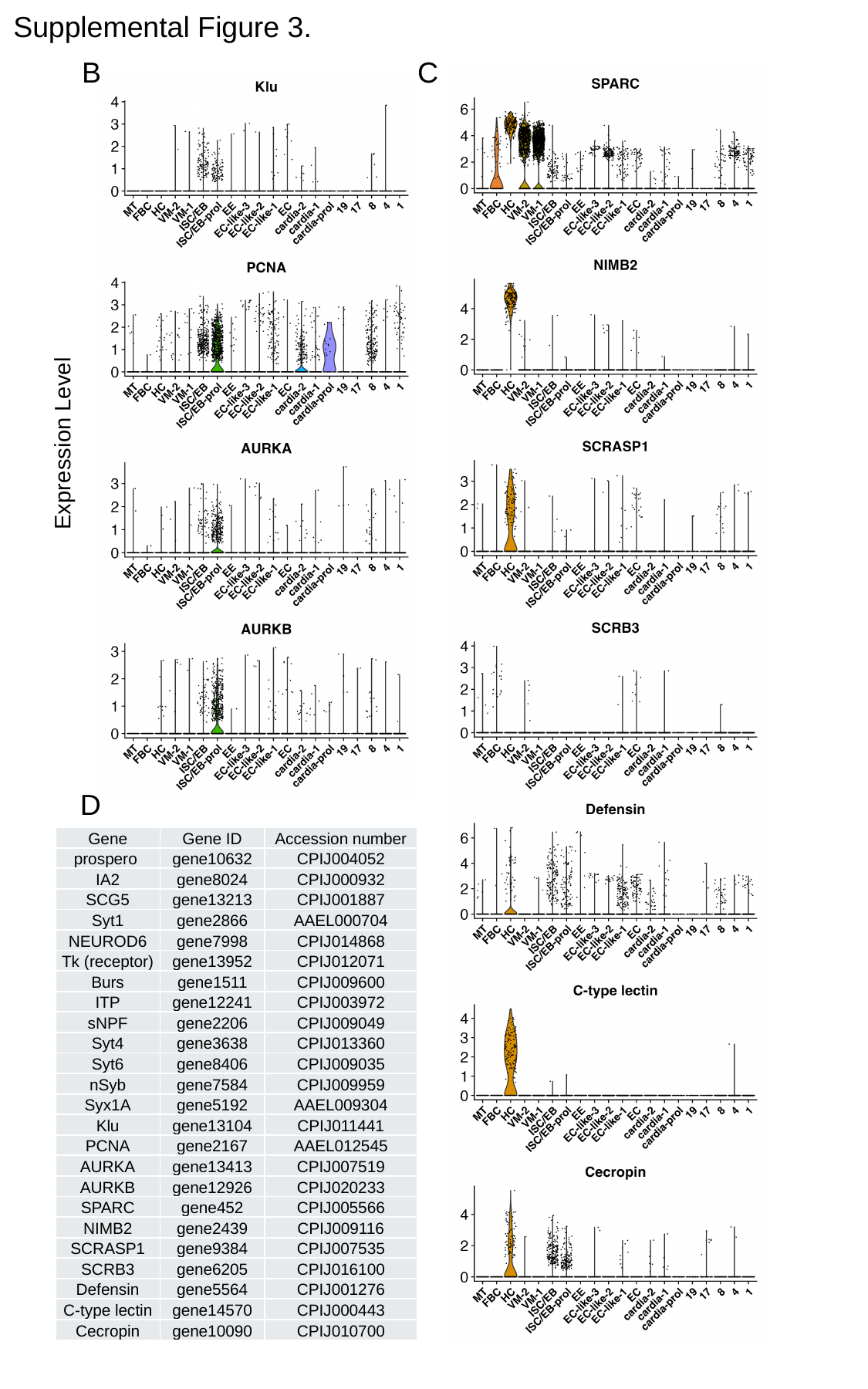

Supplemental Figure 3.
B
C
Expression Level
D
| Gene | Gene ID | Accession number |
| --- | --- | --- |
| prospero | gene10632 | CPIJ004052 |
| IA2 | gene8024 | CPIJ000932 |
| SCG5 | gene13213 | CPIJ001887 |
| Syt1 | gene2866 | AAEL000704 |
| NEUROD6 | gene7998 | CPIJ014868 |
| Tk (receptor) | gene13952 | CPIJ012071 |
| Burs | gene1511 | CPIJ009600 |
| ITP | gene12241 | CPIJ003972 |
| sNPF | gene2206 | CPIJ009049 |
| Syt4 | gene3638 | CPIJ013360 |
| Syt6 | gene8406 | CPIJ009035 |
| nSyb | gene7584 | CPIJ009959 |
| Syx1A | gene5192 | AAEL009304 |
| Klu | gene13104 | CPIJ011441 |
| PCNA | gene2167 | AAEL012545 |
| AURKA | gene13413 | CPIJ007519 |
| AURKB | gene12926 | CPIJ020233 |
| SPARC | gene452 | CPIJ005566 |
| NIMB2 | gene2439 | CPIJ009116 |
| SCRASP1 | gene9384 | CPIJ007535 |
| SCRB3 | gene6205 | CPIJ016100 |
| Defensin | gene5564 | CPIJ001276 |
| C-type lectin | gene14570 | CPIJ000443 |
| Cecropin | gene10090 | CPIJ010700 |

### Slide 5
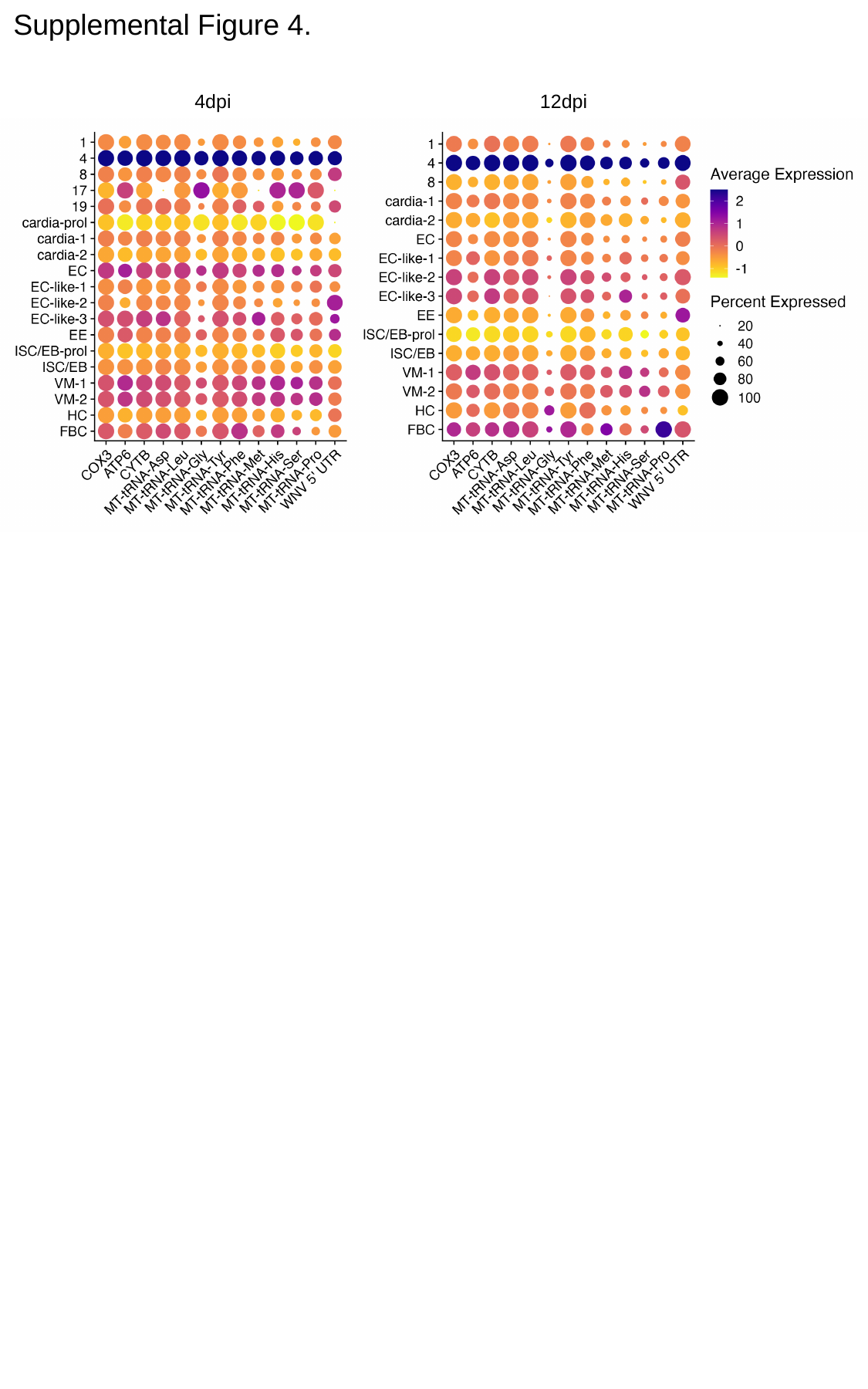

Supplemental Figure 4.
4dpi
12dpi

### Slide 6
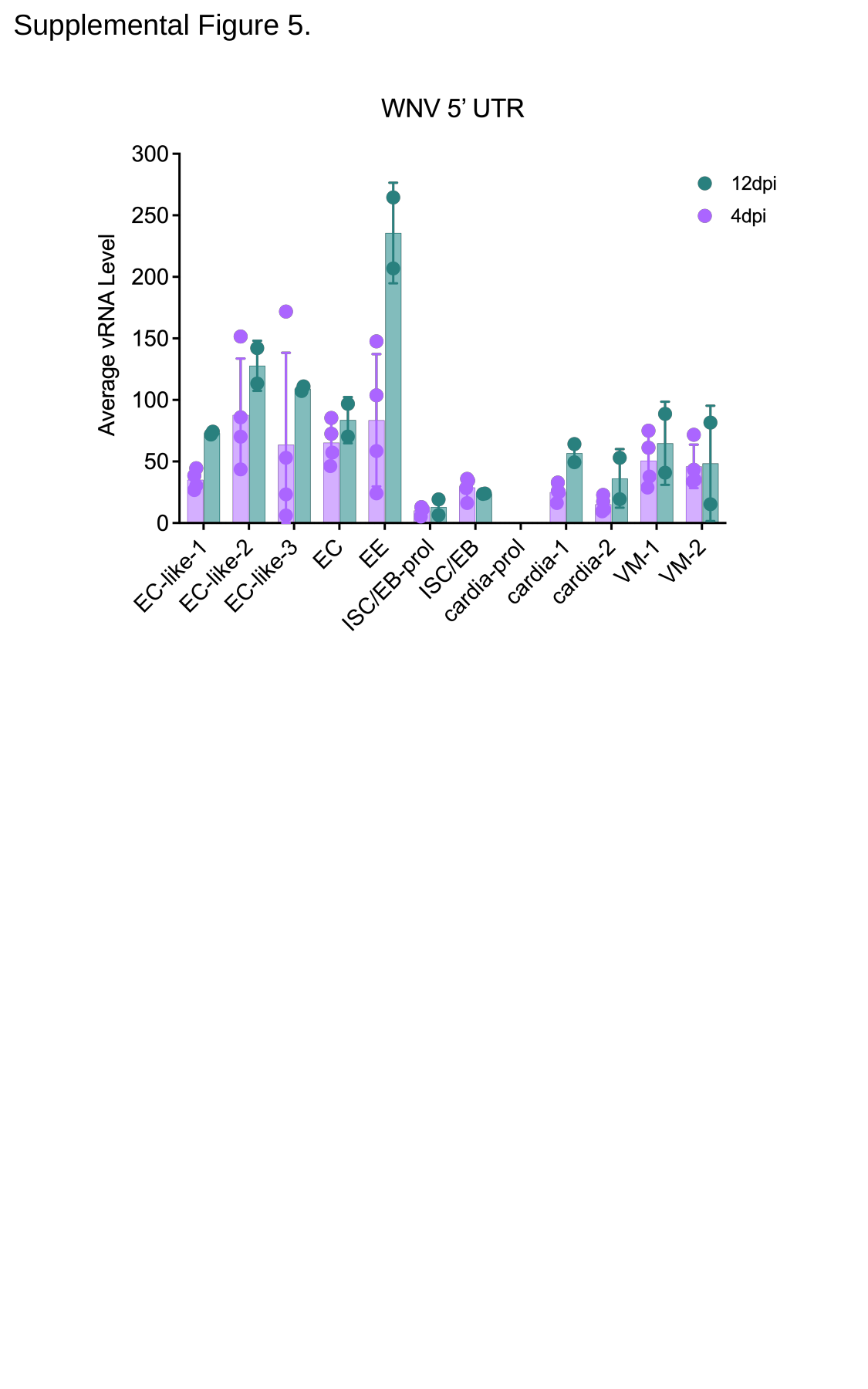

Supplemental Figure 5.

### Slide 7
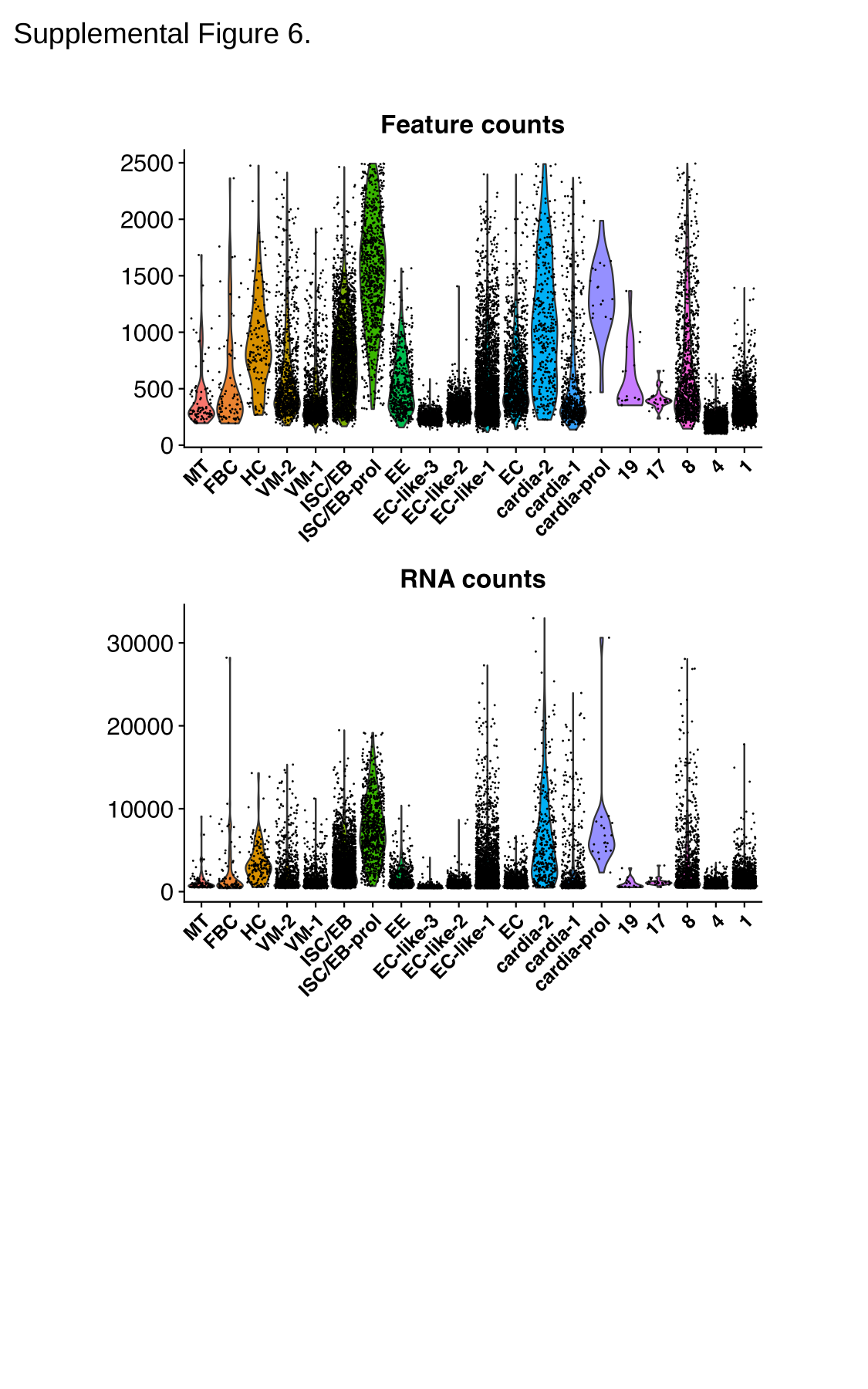

Supplemental Figure 6.

### Slide 8
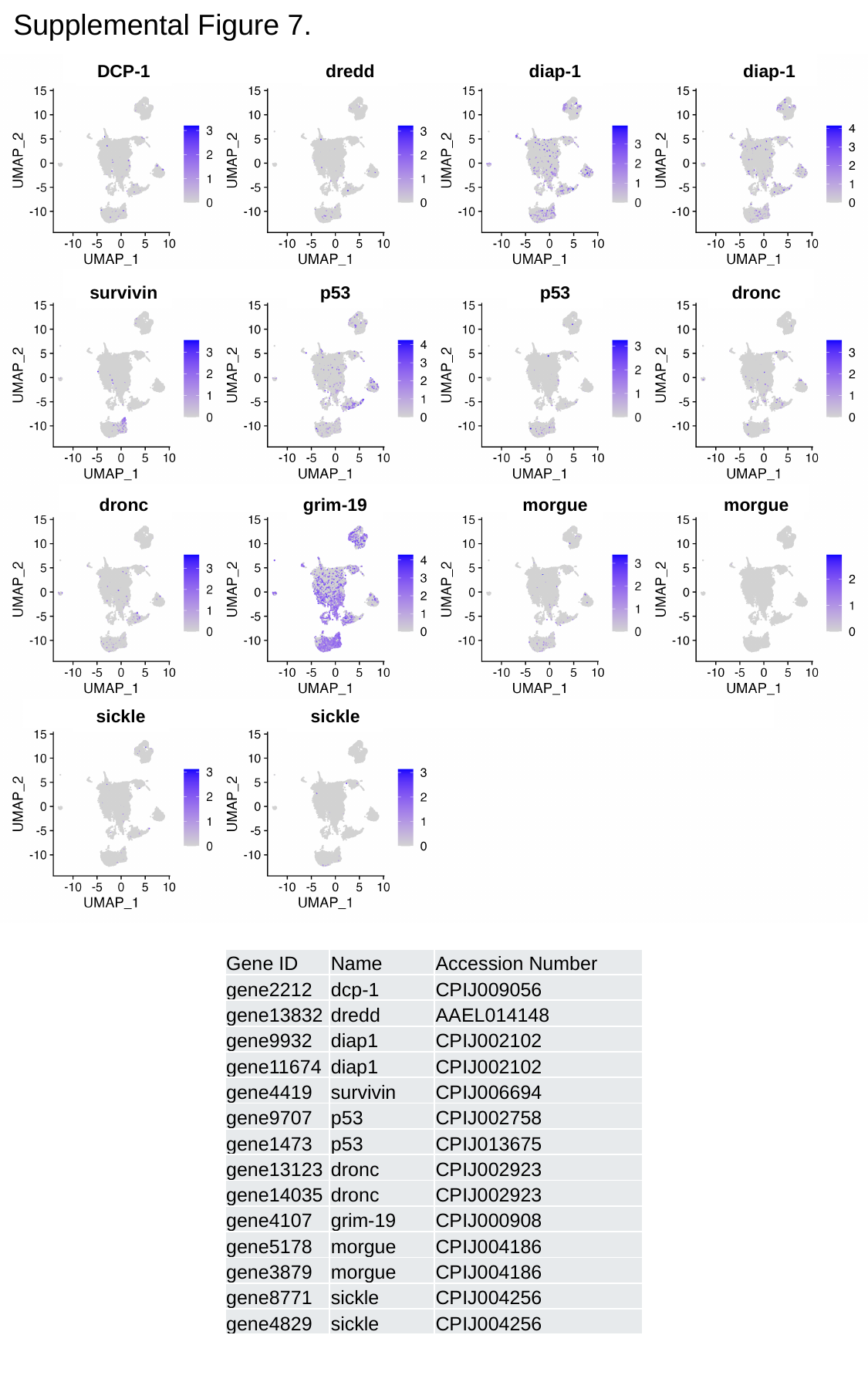

Supplemental Figure 7.
DCP-1
dredd
diap-1
diap-1
survivin
p53
p53
dronc
dronc
grim-19
morgue
morgue
sickle
sickle
| Gene ID | Name | Accession Number |
| --- | --- | --- |
| gene2212 | dcp-1 | CPIJ009056 |
| gene13832 | dredd | AAEL014148 |
| gene9932 | diap1 | CPIJ002102 |
| gene11674 | diap1 | CPIJ002102 |
| gene4419 | survivin | CPIJ006694 |
| gene9707 | p53 | CPIJ002758 |
| gene1473 | p53 | CPIJ013675 |
| gene13123 | dronc | CPIJ002923 |
| gene14035 | dronc | CPIJ002923 |
| gene4107 | grim-19 | CPIJ000908 |
| gene5178 | morgue | CPIJ004186 |
| gene3879 | morgue | CPIJ004186 |
| gene8771 | sickle | CPIJ004256 |
| gene4829 | sickle | CPIJ004256 |

### Slide 9
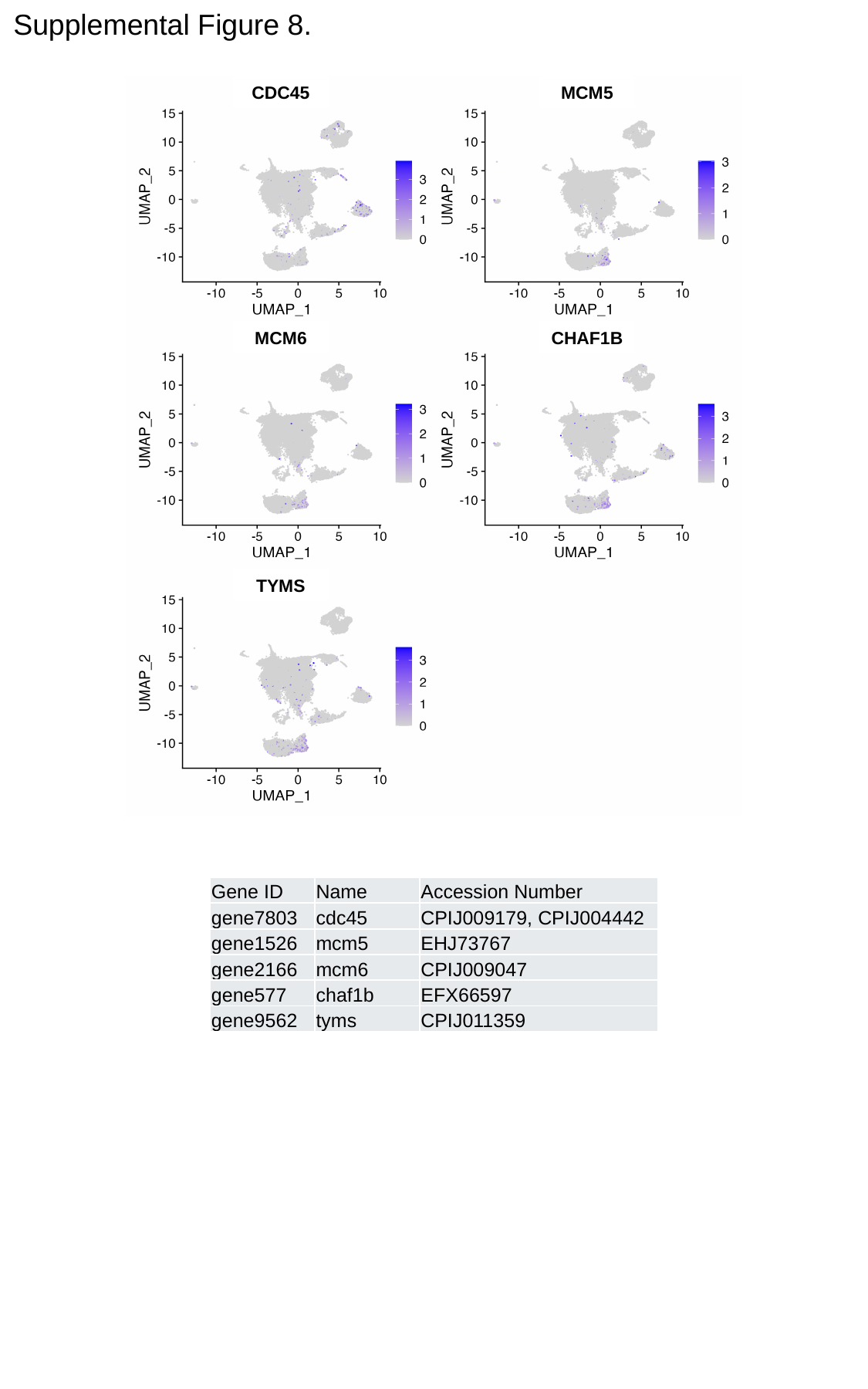

Supplemental Figure 8.
CDC45
MCM5
MCM6
CHAF1B
TYMS
| Gene ID | Name | Accession Number |
| --- | --- | --- |
| gene7803 | cdc45 | CPIJ009179, CPIJ004442 |
| gene1526 | mcm5 | EHJ73767 |
| gene2166 | mcm6 | CPIJ009047 |
| gene577 | chaf1b | EFX66597 |
| gene9562 | tyms | CPIJ011359 |

### Slide 10
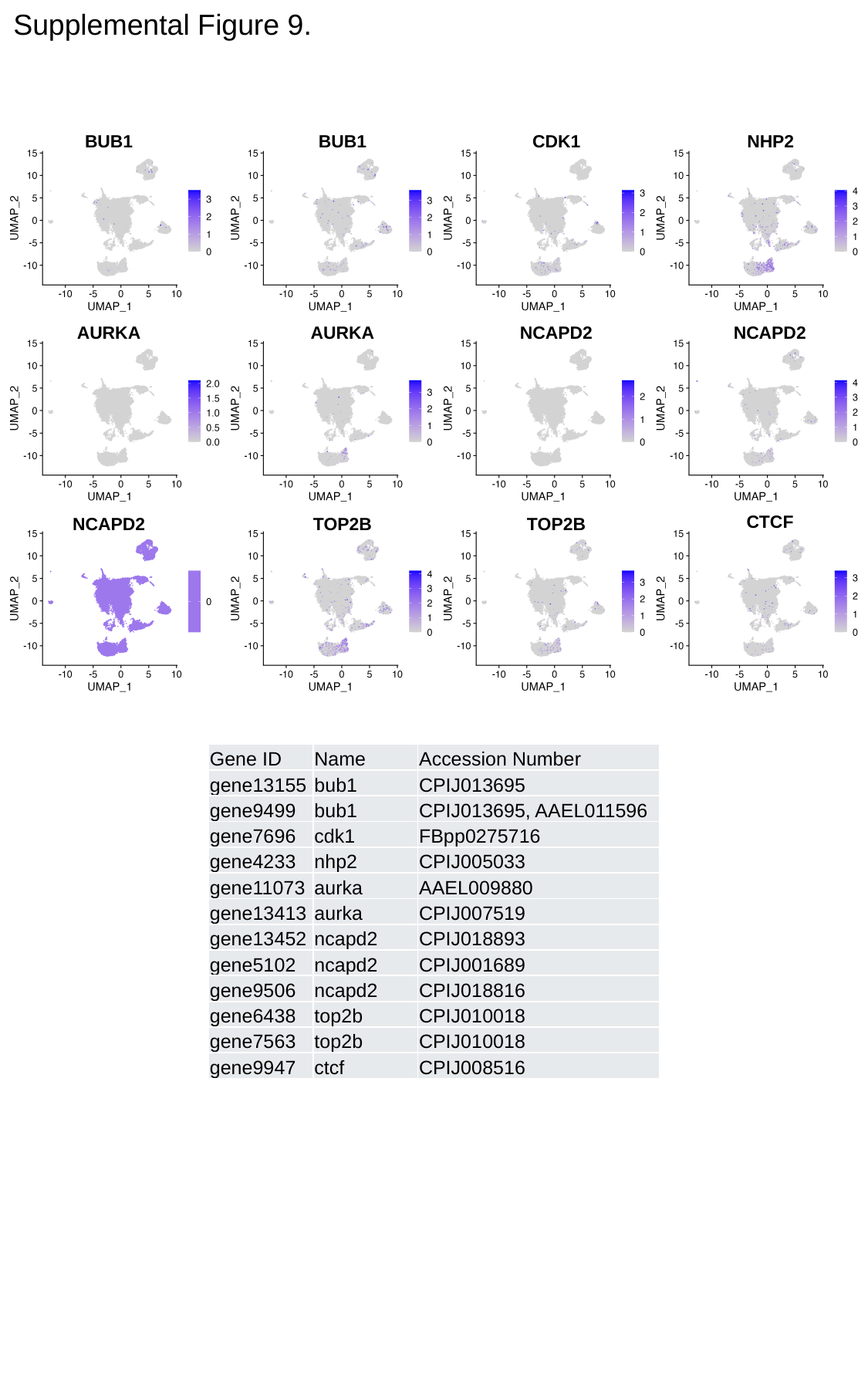

Supplemental Figure 9.
BUB1
BUB1
CDK1
NHP2
AURKA
AURKA
NCAPD2
NCAPD2
CTCF
NCAPD2
TOP2B
TOP2B
| Gene ID | Name | Accession Number |
| --- | --- | --- |
| gene13155 | bub1 | CPIJ013695 |
| gene9499 | bub1 | CPIJ013695, AAEL011596 |
| gene7696 | cdk1 | FBpp0275716 |
| gene4233 | nhp2 | CPIJ005033 |
| gene11073 | aurka | AAEL009880 |
| gene13413 | aurka | CPIJ007519 |
| gene13452 | ncapd2 | CPIJ018893 |
| gene5102 | ncapd2 | CPIJ001689 |
| gene9506 | ncapd2 | CPIJ018816 |
| gene6438 | top2b | CPIJ010018 |
| gene7563 | top2b | CPIJ010018 |
| gene9947 | ctcf | CPIJ008516 |

### Slide 11
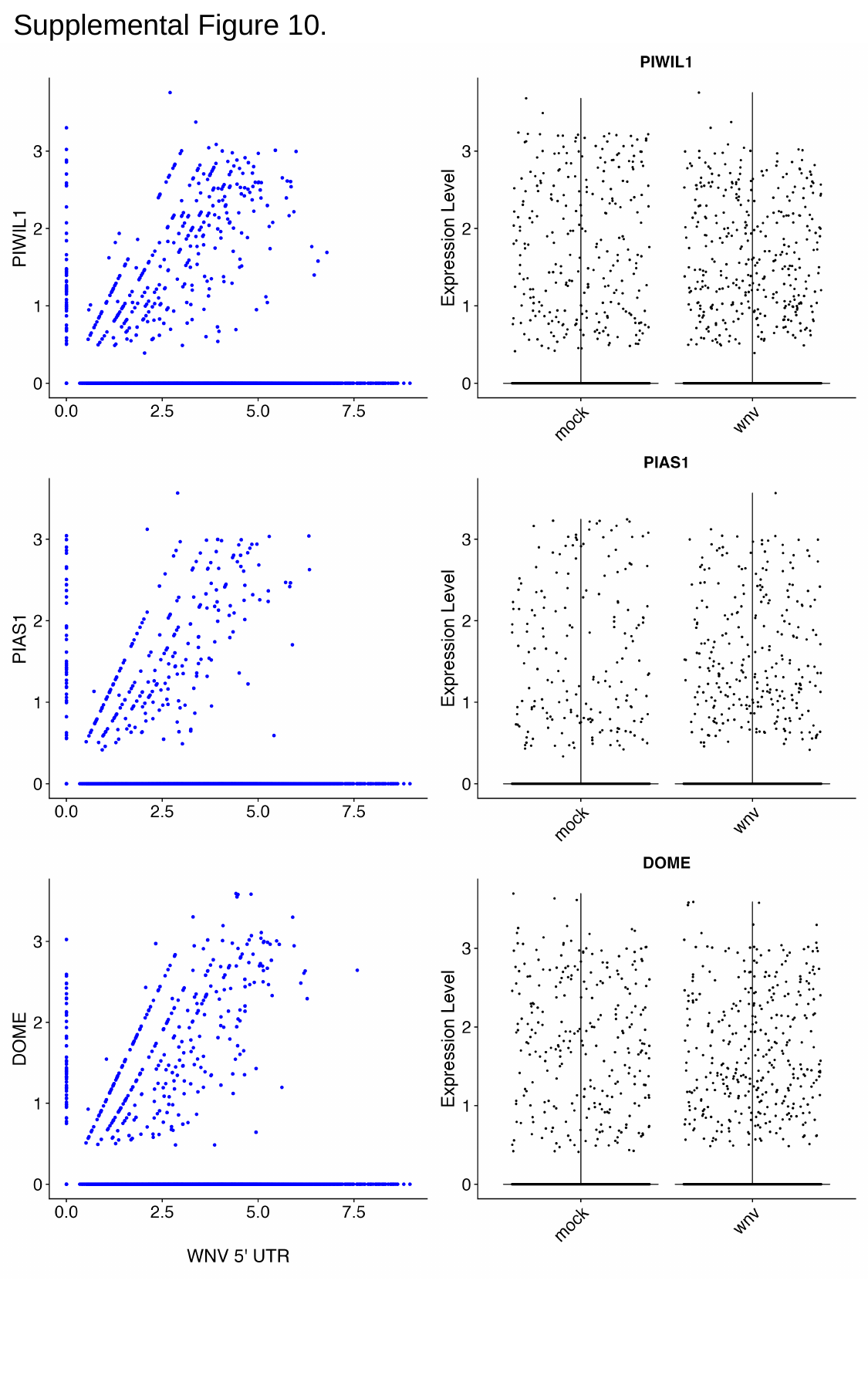

Supplemental Figure 10.

### Slide 12
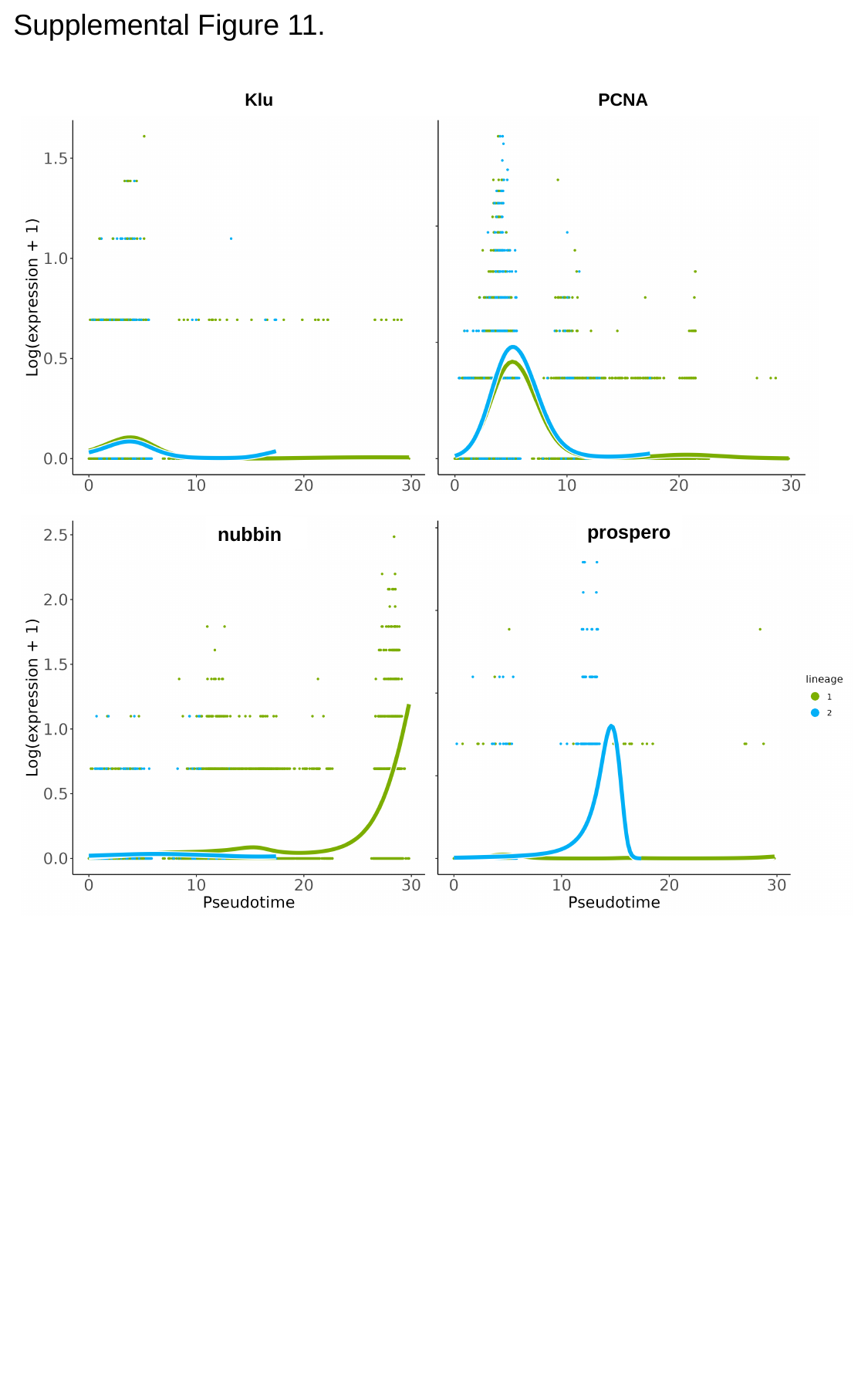

Supplemental Figure 11.
Klu
PCNA
prospero
nubbin
POU2F1
PROX1

### Slide 13
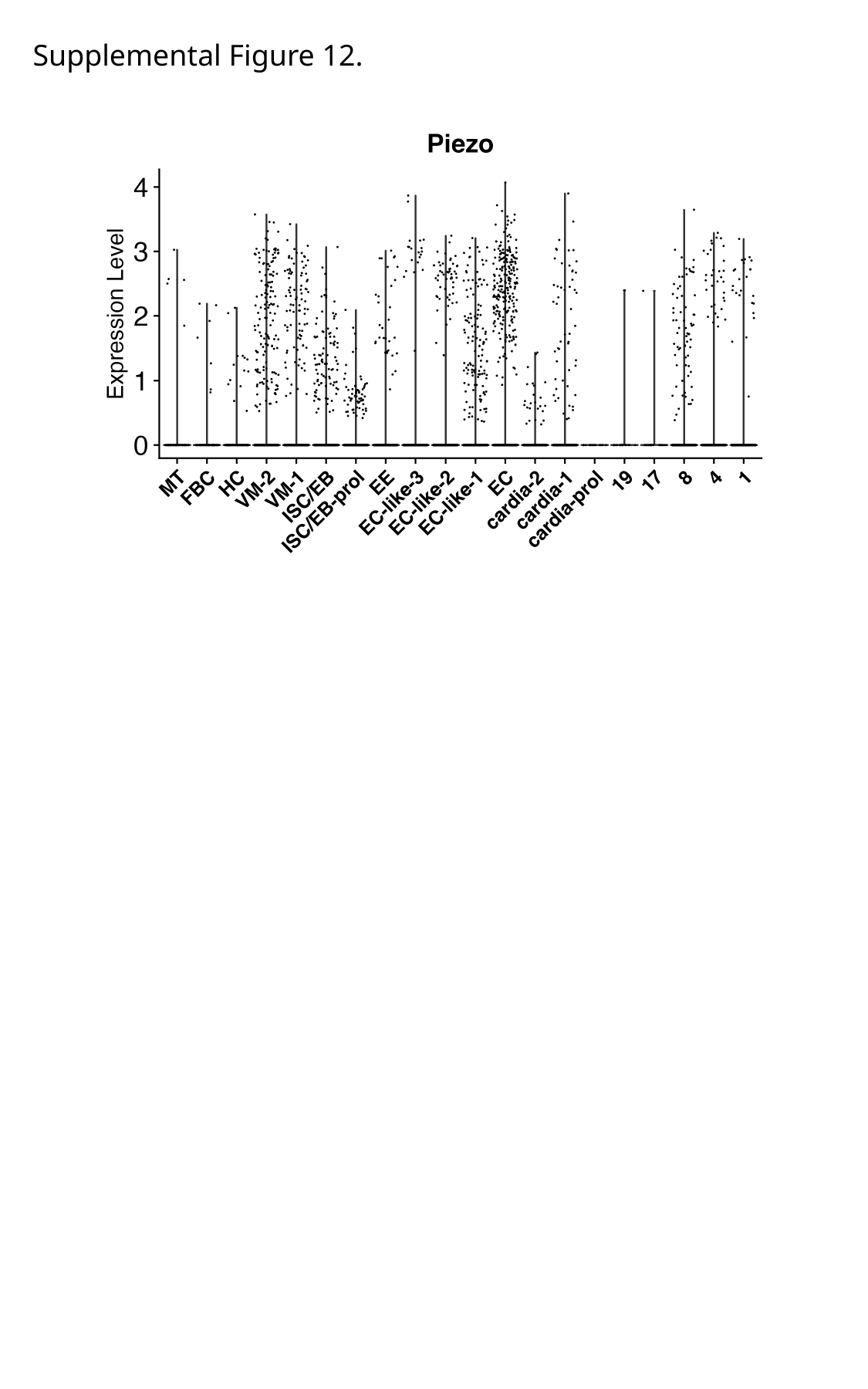

Supplemental Figure 12.

### Slide 14
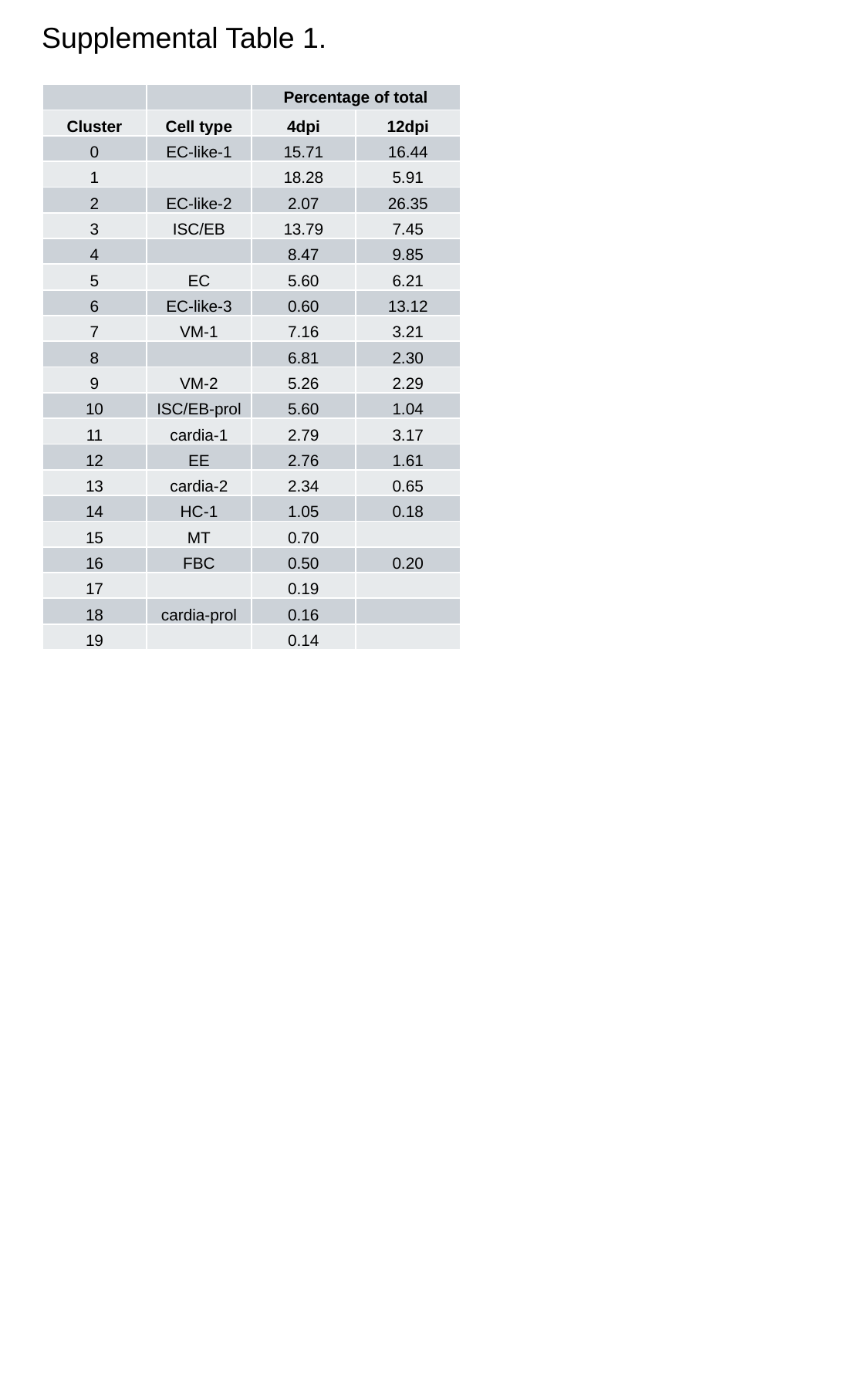

Supplemental Table 1.
| | | Percentage of total | |
| --- | --- | --- | --- |
| Cluster | Cell type | 4dpi | 12dpi |
| 0 | EC-like-1 | 15.71 | 16.44 |
| 1 | | 18.28 | 5.91 |
| 2 | EC-like-2 | 2.07 | 26.35 |
| 3 | ISC/EB | 13.79 | 7.45 |
| 4 | | 8.47 | 9.85 |
| 5 | EC | 5.60 | 6.21 |
| 6 | EC-like-3 | 0.60 | 13.12 |
| 7 | VM-1 | 7.16 | 3.21 |
| 8 | | 6.81 | 2.30 |
| 9 | VM-2 | 5.26 | 2.29 |
| 10 | ISC/EB-prol | 5.60 | 1.04 |
| 11 | cardia-1 | 2.79 | 3.17 |
| 12 | EE | 2.76 | 1.61 |
| 13 | cardia-2 | 2.34 | 0.65 |
| 14 | HC-1 | 1.05 | 0.18 |
| 15 | MT | 0.70 | |
| 16 | FBC | 0.50 | 0.20 |
| 17 | | 0.19 | |
| 18 | cardia-prol | 0.16 | |
| 19 | | 0.14 | |

### Slide 15
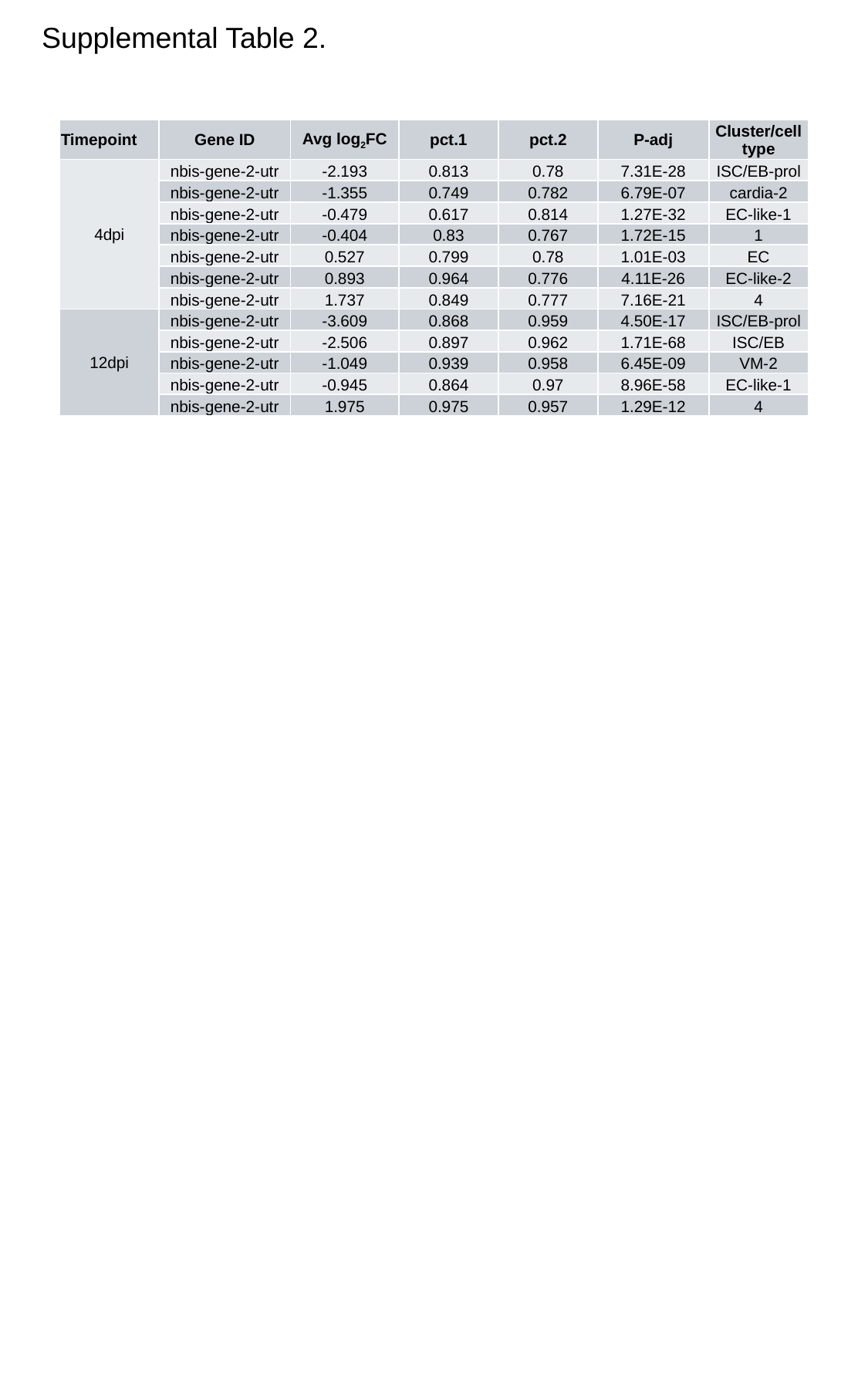

Supplemental Table 2.
| Timepoint | Gene ID | Avg log_2_FC | pct.1 | pct.2 | P-adj | Cluster/cell type |
| --- | --- | --- | --- | --- | --- | --- |
| 4dpi | nbis-gene-2-utr | -2.193 | 0.813 | 0.78 | 7.31E-28 | ISC/EB-prol |
| | nbis-gene-2-utr | -1.355 | 0.749 | 0.782 | 6.79E-07 | cardia-2 |
| | nbis-gene-2-utr | -0.479 | 0.617 | 0.814 | 1.27E-32 | EC-like-1 |
| | nbis-gene-2-utr | -0.404 | 0.83 | 0.767 | 1.72E-15 | 1 |
| | nbis-gene-2-utr | 0.527 | 0.799 | 0.78 | 1.01E-03 | EC |
| | nbis-gene-2-utr | 0.893 | 0.964 | 0.776 | 4.11E-26 | EC-like-2 |
| | nbis-gene-2-utr | 1.737 | 0.849 | 0.777 | 7.16E-21 | 4 |
| 12dpi | nbis-gene-2-utr | -3.609 | 0.868 | 0.959 | 4.50E-17 | ISC/EB-prol |
| | nbis-gene-2-utr | -2.506 | 0.897 | 0.962 | 1.71E-68 | ISC/EB |
| | nbis-gene-2-utr | -1.049 | 0.939 | 0.958 | 6.45E-09 | VM-2 |
| | nbis-gene-2-utr | -0.945 | 0.864 | 0.97 | 8.96E-58 | EC-like-1 |
| | nbis-gene-2-utr | 1.975 | 0.975 | 0.957 | 1.29E-12 | 4 |

### Slide 16
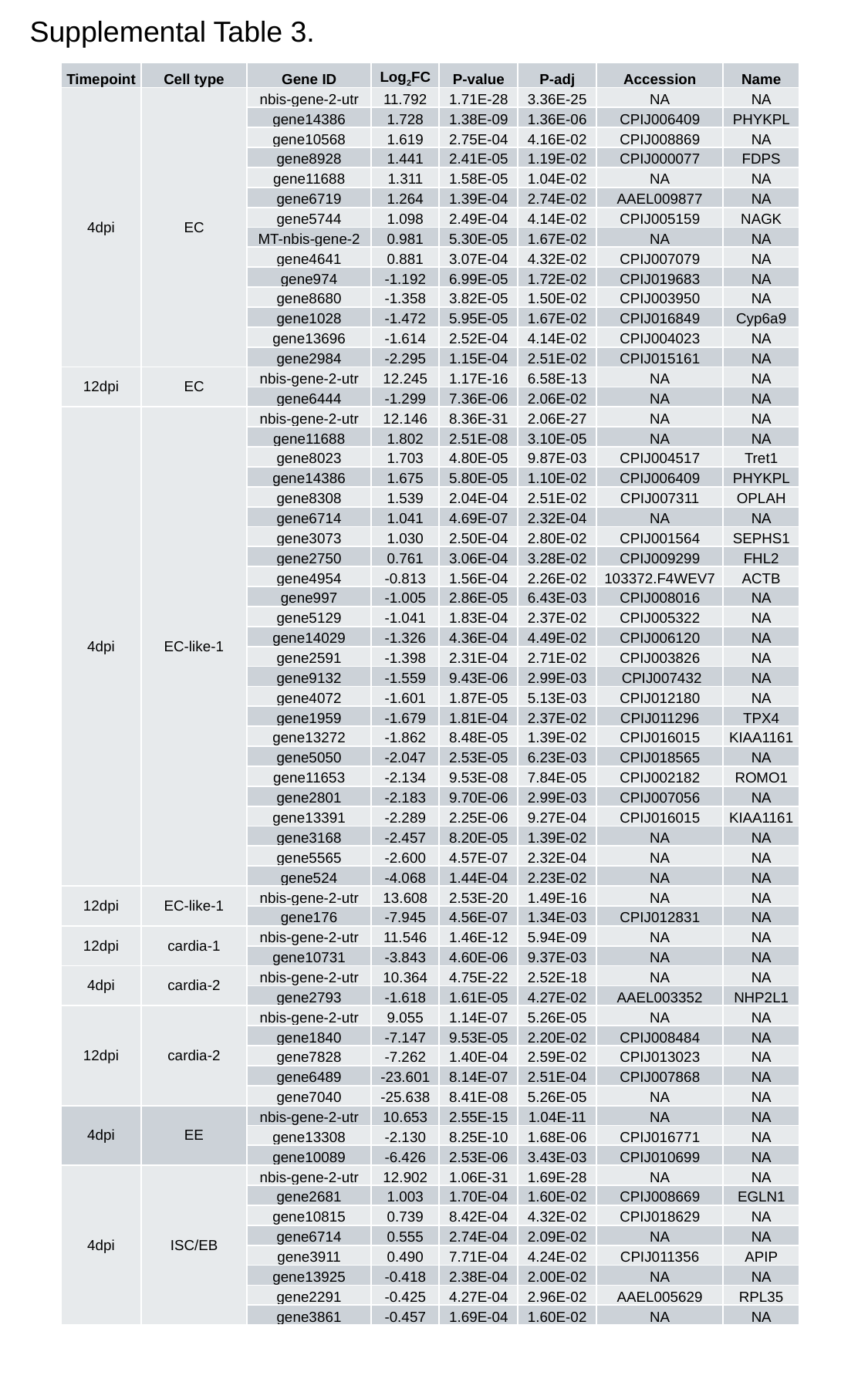

Supplemental Table 3.
| Timepoint | Cell type | Gene ID | Log_2_FC | P-value | P-adj | Accession | Name |
| --- | --- | --- | --- | --- | --- | --- | --- |
| 4dpi | EC | nbis-gene-2-utr | 11.792 | 1.71E-28 | 3.36E-25 | NA | NA |
| | | gene14386 | 1.728 | 1.38E-09 | 1.36E-06 | CPIJ006409 | PHYKPL |
| | | gene10568 | 1.619 | 2.75E-04 | 4.16E-02 | CPIJ008869 | NA |
| | | gene8928 | 1.441 | 2.41E-05 | 1.19E-02 | CPIJ000077 | FDPS |
| | | gene11688 | 1.311 | 1.58E-05 | 1.04E-02 | NA | NA |
| | | gene6719 | 1.264 | 1.39E-04 | 2.74E-02 | AAEL009877 | NA |
| | | gene5744 | 1.098 | 2.49E-04 | 4.14E-02 | CPIJ005159 | NAGK |
| | | MT-nbis-gene-2 | 0.981 | 5.30E-05 | 1.67E-02 | NA | NA |
| | | gene4641 | 0.881 | 3.07E-04 | 4.32E-02 | CPIJ007079 | NA |
| | | gene974 | -1.192 | 6.99E-05 | 1.72E-02 | CPIJ019683 | NA |
| | | gene8680 | -1.358 | 3.82E-05 | 1.50E-02 | CPIJ003950 | NA |
| | | gene1028 | -1.472 | 5.95E-05 | 1.67E-02 | CPIJ016849 | Cyp6a9 |
| | | gene13696 | -1.614 | 2.52E-04 | 4.14E-02 | CPIJ004023 | NA |
| | | gene2984 | -2.295 | 1.15E-04 | 2.51E-02 | CPIJ015161 | NA |
| 12dpi | EC | nbis-gene-2-utr | 12.245 | 1.17E-16 | 6.58E-13 | NA | NA |
| | | gene6444 | -1.299 | 7.36E-06 | 2.06E-02 | NA | NA |
| 4dpi | EC-like-1 | nbis-gene-2-utr | 12.146 | 8.36E-31 | 2.06E-27 | NA | NA |
| | | gene11688 | 1.802 | 2.51E-08 | 3.10E-05 | NA | NA |
| | | gene8023 | 1.703 | 4.80E-05 | 9.87E-03 | CPIJ004517 | Tret1 |
| | | gene14386 | 1.675 | 5.80E-05 | 1.10E-02 | CPIJ006409 | PHYKPL |
| | | gene8308 | 1.539 | 2.04E-04 | 2.51E-02 | CPIJ007311 | OPLAH |
| | | gene6714 | 1.041 | 4.69E-07 | 2.32E-04 | NA | NA |
| | | gene3073 | 1.030 | 2.50E-04 | 2.80E-02 | CPIJ001564 | SEPHS1 |
| | | gene2750 | 0.761 | 3.06E-04 | 3.28E-02 | CPIJ009299 | FHL2 |
| | | gene4954 | -0.813 | 1.56E-04 | 2.26E-02 | 103372.F4WEV7 | ACTB |
| | | gene997 | -1.005 | 2.86E-05 | 6.43E-03 | CPIJ008016 | NA |
| | | gene5129 | -1.041 | 1.83E-04 | 2.37E-02 | CPIJ005322 | NA |
| | | gene14029 | -1.326 | 4.36E-04 | 4.49E-02 | CPIJ006120 | NA |
| | | gene2591 | -1.398 | 2.31E-04 | 2.71E-02 | CPIJ003826 | NA |
| | | gene9132 | -1.559 | 9.43E-06 | 2.99E-03 | CPIJ007432 | NA |
| | | gene4072 | -1.601 | 1.87E-05 | 5.13E-03 | CPIJ012180 | NA |
| | | gene1959 | -1.679 | 1.81E-04 | 2.37E-02 | CPIJ011296 | TPX4 |
| | | gene13272 | -1.862 | 8.48E-05 | 1.39E-02 | CPIJ016015 | KIAA1161 |
| | | gene5050 | -2.047 | 2.53E-05 | 6.23E-03 | CPIJ018565 | NA |
| | | gene11653 | -2.134 | 9.53E-08 | 7.84E-05 | CPIJ002182 | ROMO1 |
| | | gene2801 | -2.183 | 9.70E-06 | 2.99E-03 | CPIJ007056 | NA |
| | | gene13391 | -2.289 | 2.25E-06 | 9.27E-04 | CPIJ016015 | KIAA1161 |
| | | gene3168 | -2.457 | 8.20E-05 | 1.39E-02 | NA | NA |
| | | gene5565 | -2.600 | 4.57E-07 | 2.32E-04 | NA | NA |
| | | gene524 | -4.068 | 1.44E-04 | 2.23E-02 | NA | NA |
| 12dpi | EC-like-1 | nbis-gene-2-utr | 13.608 | 2.53E-20 | 1.49E-16 | NA | NA |
| | | gene176 | -7.945 | 4.56E-07 | 1.34E-03 | CPIJ012831 | NA |
| 12dpi | cardia-1 | nbis-gene-2-utr | 11.546 | 1.46E-12 | 5.94E-09 | NA | NA |
| | | gene10731 | -3.843 | 4.60E-06 | 9.37E-03 | NA | NA |
| 4dpi | cardia-2 | nbis-gene-2-utr | 10.364 | 4.75E-22 | 2.52E-18 | NA | NA |
| | | gene2793 | -1.618 | 1.61E-05 | 4.27E-02 | AAEL003352 | NHP2L1 |
| 12dpi | cardia-2 | nbis-gene-2-utr | 9.055 | 1.14E-07 | 5.26E-05 | NA | NA |
| | | gene1840 | -7.147 | 9.53E-05 | 2.20E-02 | CPIJ008484 | NA |
| | | gene7828 | -7.262 | 1.40E-04 | 2.59E-02 | CPIJ013023 | NA |
| | | gene6489 | -23.601 | 8.14E-07 | 2.51E-04 | CPIJ007868 | NA |
| | | gene7040 | -25.638 | 8.41E-08 | 5.26E-05 | NA | NA |
| 4dpi | EE | nbis-gene-2-utr | 10.653 | 2.55E-15 | 1.04E-11 | NA | NA |
| | | gene13308 | -2.130 | 8.25E-10 | 1.68E-06 | CPIJ016771 | NA |
| | | gene10089 | -6.426 | 2.53E-06 | 3.43E-03 | CPIJ010699 | NA |
| 4dpi | ISC/EB | nbis-gene-2-utr | 12.902 | 1.06E-31 | 1.69E-28 | NA | NA |
| | | gene2681 | 1.003 | 1.70E-04 | 1.60E-02 | CPIJ008669 | EGLN1 |
| | | gene10815 | 0.739 | 8.42E-04 | 4.32E-02 | CPIJ018629 | NA |
| | | gene6714 | 0.555 | 2.74E-04 | 2.09E-02 | NA | NA |
| | | gene3911 | 0.490 | 7.71E-04 | 4.24E-02 | CPIJ011356 | APIP |
| | | gene13925 | -0.418 | 2.38E-04 | 2.00E-02 | NA | NA |
| | | gene2291 | -0.425 | 4.27E-04 | 2.96E-02 | AAEL005629 | RPL35 |
| | | gene3861 | -0.457 | 1.69E-04 | 1.60E-02 | NA | NA |

### Slide 17
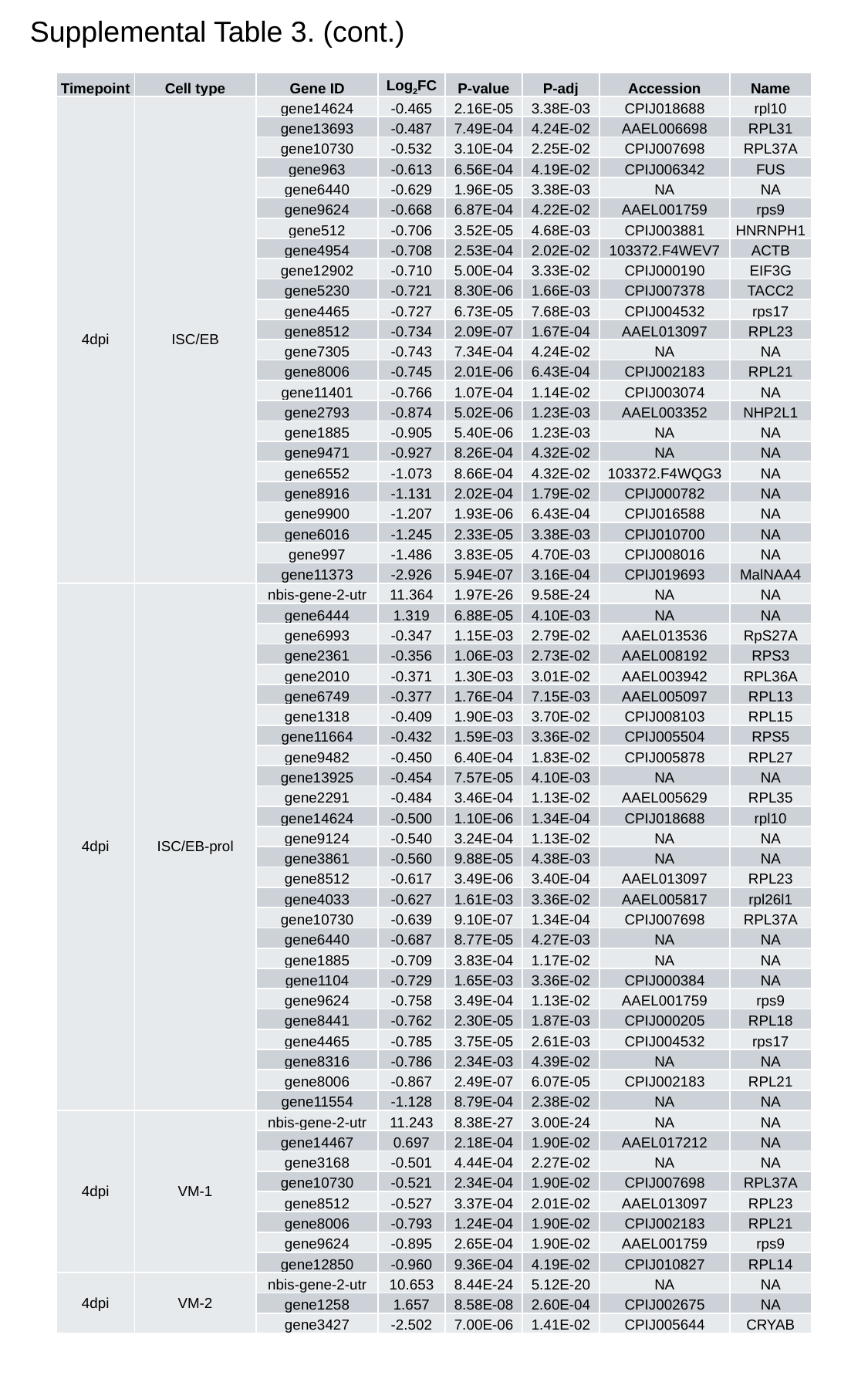

Supplemental Table 3. (cont.)
| Timepoint | Cell type | Gene ID | Log_2_FC | P-value | P-adj | Accession | Name |
| --- | --- | --- | --- | --- | --- | --- | --- |
| 4dpi | ISC/EB | gene14624 | -0.465 | 2.16E-05 | 3.38E-03 | CPIJ018688 | rpl10 |
| | | gene13693 | -0.487 | 7.49E-04 | 4.24E-02 | AAEL006698 | RPL31 |
| | | gene10730 | -0.532 | 3.10E-04 | 2.25E-02 | CPIJ007698 | RPL37A |
| | | gene963 | -0.613 | 6.56E-04 | 4.19E-02 | CPIJ006342 | FUS |
| | | gene6440 | -0.629 | 1.96E-05 | 3.38E-03 | NA | NA |
| | | gene9624 | -0.668 | 6.87E-04 | 4.22E-02 | AAEL001759 | rps9 |
| | | gene512 | -0.706 | 3.52E-05 | 4.68E-03 | CPIJ003881 | HNRNPH1 |
| | | gene4954 | -0.708 | 2.53E-04 | 2.02E-02 | 103372.F4WEV7 | ACTB |
| | | gene12902 | -0.710 | 5.00E-04 | 3.33E-02 | CPIJ000190 | EIF3G |
| | | gene5230 | -0.721 | 8.30E-06 | 1.66E-03 | CPIJ007378 | TACC2 |
| | | gene4465 | -0.727 | 6.73E-05 | 7.68E-03 | CPIJ004532 | rps17 |
| | | gene8512 | -0.734 | 2.09E-07 | 1.67E-04 | AAEL013097 | RPL23 |
| | | gene7305 | -0.743 | 7.34E-04 | 4.24E-02 | NA | NA |
| | | gene8006 | -0.745 | 2.01E-06 | 6.43E-04 | CPIJ002183 | RPL21 |
| | | gene11401 | -0.766 | 1.07E-04 | 1.14E-02 | CPIJ003074 | NA |
| | | gene2793 | -0.874 | 5.02E-06 | 1.23E-03 | AAEL003352 | NHP2L1 |
| | | gene1885 | -0.905 | 5.40E-06 | 1.23E-03 | NA | NA |
| | | gene9471 | -0.927 | 8.26E-04 | 4.32E-02 | NA | NA |
| | | gene6552 | -1.073 | 8.66E-04 | 4.32E-02 | 103372.F4WQG3 | NA |
| | | gene8916 | -1.131 | 2.02E-04 | 1.79E-02 | CPIJ000782 | NA |
| | | gene9900 | -1.207 | 1.93E-06 | 6.43E-04 | CPIJ016588 | NA |
| | | gene6016 | -1.245 | 2.33E-05 | 3.38E-03 | CPIJ010700 | NA |
| | | gene997 | -1.486 | 3.83E-05 | 4.70E-03 | CPIJ008016 | NA |
| | | gene11373 | -2.926 | 5.94E-07 | 3.16E-04 | CPIJ019693 | MalNAA4 |
| 4dpi | ISC/EB-prol | nbis-gene-2-utr | 11.364 | 1.97E-26 | 9.58E-24 | NA | NA |
| | | gene6444 | 1.319 | 6.88E-05 | 4.10E-03 | NA | NA |
| | | gene6993 | -0.347 | 1.15E-03 | 2.79E-02 | AAEL013536 | RpS27A |
| | | gene2361 | -0.356 | 1.06E-03 | 2.73E-02 | AAEL008192 | RPS3 |
| | | gene2010 | -0.371 | 1.30E-03 | 3.01E-02 | AAEL003942 | RPL36A |
| | | gene6749 | -0.377 | 1.76E-04 | 7.15E-03 | AAEL005097 | RPL13 |
| | | gene1318 | -0.409 | 1.90E-03 | 3.70E-02 | CPIJ008103 | RPL15 |
| | | gene11664 | -0.432 | 1.59E-03 | 3.36E-02 | CPIJ005504 | RPS5 |
| | | gene9482 | -0.450 | 6.40E-04 | 1.83E-02 | CPIJ005878 | RPL27 |
| | | gene13925 | -0.454 | 7.57E-05 | 4.10E-03 | NA | NA |
| | | gene2291 | -0.484 | 3.46E-04 | 1.13E-02 | AAEL005629 | RPL35 |
| | | gene14624 | -0.500 | 1.10E-06 | 1.34E-04 | CPIJ018688 | rpl10 |
| | | gene9124 | -0.540 | 3.24E-04 | 1.13E-02 | NA | NA |
| | | gene3861 | -0.560 | 9.88E-05 | 4.38E-03 | NA | NA |
| | | gene8512 | -0.617 | 3.49E-06 | 3.40E-04 | AAEL013097 | RPL23 |
| | | gene4033 | -0.627 | 1.61E-03 | 3.36E-02 | AAEL005817 | rpl26l1 |
| | | gene10730 | -0.639 | 9.10E-07 | 1.34E-04 | CPIJ007698 | RPL37A |
| | | gene6440 | -0.687 | 8.77E-05 | 4.27E-03 | NA | NA |
| | | gene1885 | -0.709 | 3.83E-04 | 1.17E-02 | NA | NA |
| | | gene1104 | -0.729 | 1.65E-03 | 3.36E-02 | CPIJ000384 | NA |
| | | gene9624 | -0.758 | 3.49E-04 | 1.13E-02 | AAEL001759 | rps9 |
| | | gene8441 | -0.762 | 2.30E-05 | 1.87E-03 | CPIJ000205 | RPL18 |
| | | gene4465 | -0.785 | 3.75E-05 | 2.61E-03 | CPIJ004532 | rps17 |
| | | gene8316 | -0.786 | 2.34E-03 | 4.39E-02 | NA | NA |
| | | gene8006 | -0.867 | 2.49E-07 | 6.07E-05 | CPIJ002183 | RPL21 |
| | | gene11554 | -1.128 | 8.79E-04 | 2.38E-02 | NA | NA |
| 4dpi | VM-1 | nbis-gene-2-utr | 11.243 | 8.38E-27 | 3.00E-24 | NA | NA |
| | | gene14467 | 0.697 | 2.18E-04 | 1.90E-02 | AAEL017212 | NA |
| | | gene3168 | -0.501 | 4.44E-04 | 2.27E-02 | NA | NA |
| | | gene10730 | -0.521 | 2.34E-04 | 1.90E-02 | CPIJ007698 | RPL37A |
| | | gene8512 | -0.527 | 3.37E-04 | 2.01E-02 | AAEL013097 | RPL23 |
| | | gene8006 | -0.793 | 1.24E-04 | 1.90E-02 | CPIJ002183 | RPL21 |
| | | gene9624 | -0.895 | 2.65E-04 | 1.90E-02 | AAEL001759 | rps9 |
| | | gene12850 | -0.960 | 9.36E-04 | 4.19E-02 | CPIJ010827 | RPL14 |
| 4dpi | VM-2 | nbis-gene-2-utr | 10.653 | 8.44E-24 | 5.12E-20 | NA | NA |
| | | gene1258 | 1.657 | 8.58E-08 | 2.60E-04 | CPIJ002675 | NA |
| | | gene3427 | -2.502 | 7.00E-06 | 1.41E-02 | CPIJ005644 | CRYAB |

### Slide 18
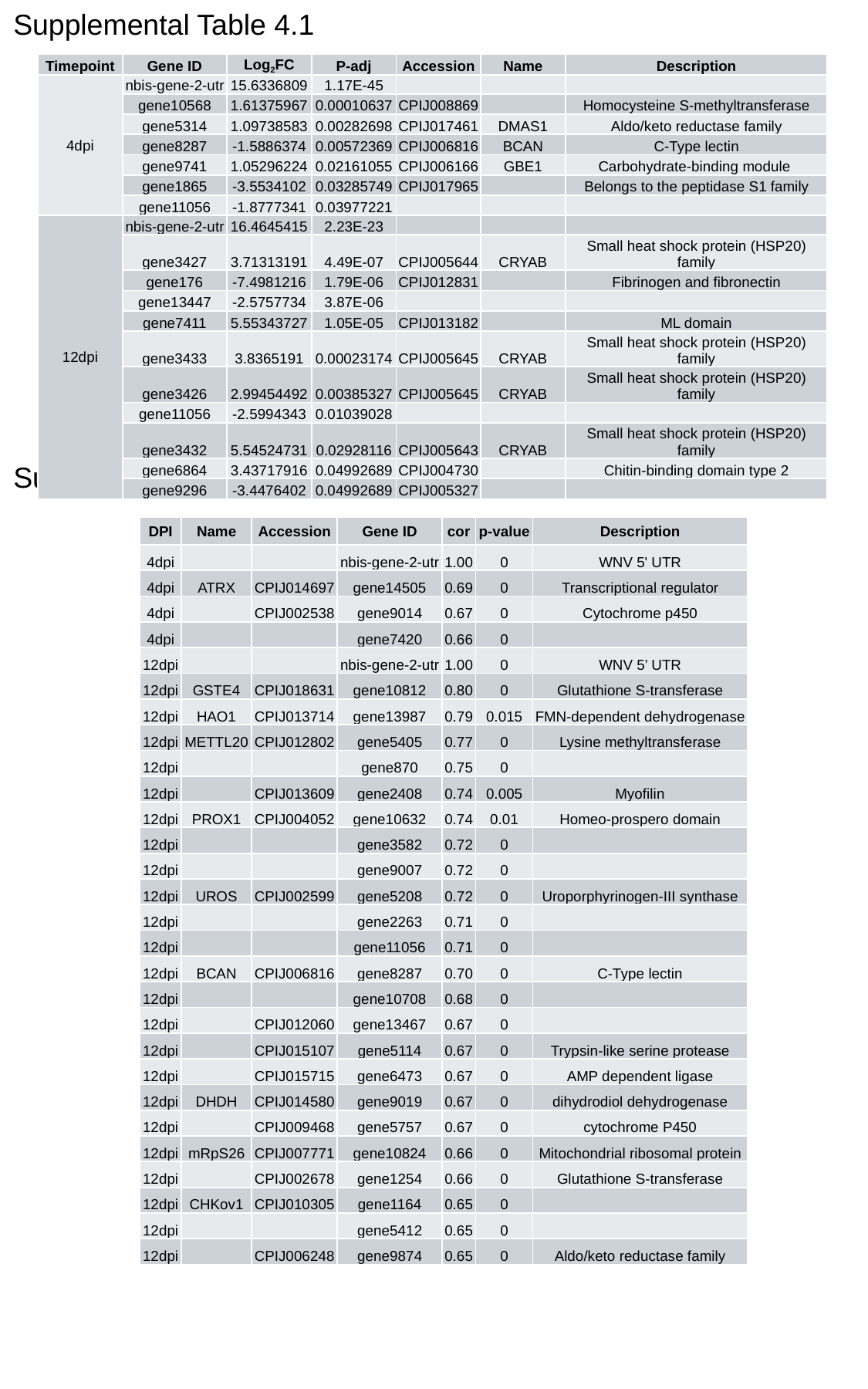

Supplemental Table 4.1
| Timepoint | Gene ID | Log_2_FC | P-adj | Accession | Name | Description |
| --- | --- | --- | --- | --- | --- | --- |
| 4dpi | nbis-gene-2-utr | 15.6336809 | 1.17E-45 | | | |
| | gene10568 | 1.61375967 | 0.00010637 | CPIJ008869 | | Homocysteine S-methyltransferase |
| | gene5314 | 1.09738583 | 0.00282698 | CPIJ017461 | DMAS1 | Aldo/keto reductase family |
| | gene8287 | -1.5886374 | 0.00572369 | CPIJ006816 | BCAN | C-Type lectin |
| | gene9741 | 1.05296224 | 0.02161055 | CPIJ006166 | GBE1 | Carbohydrate-binding module |
| | gene1865 | -3.5534102 | 0.03285749 | CPIJ017965 | | Belongs to the peptidase S1 family |
| | gene11056 | -1.8777341 | 0.03977221 | | | |
| 12dpi | nbis-gene-2-utr | 16.4645415 | 2.23E-23 | | | |
| | gene3427 | 3.71313191 | 4.49E-07 | CPIJ005644 | CRYAB | Small heat shock protein (HSP20) family |
| | gene176 | -7.4981216 | 1.79E-06 | CPIJ012831 | | Fibrinogen and fibronectin |
| | gene13447 | -2.5757734 | 3.87E-06 | | | |
| | gene7411 | 5.55343727 | 1.05E-05 | CPIJ013182 | | ML domain |
| | gene3433 | 3.8365191 | 0.00023174 | CPIJ005645 | CRYAB | Small heat shock protein (HSP20) family |
| | gene3426 | 2.99454492 | 0.00385327 | CPIJ005645 | CRYAB | Small heat shock protein (HSP20) family |
| | gene11056 | -2.5994343 | 0.01039028 | | | |
| | gene3432 | 5.54524731 | 0.02928116 | CPIJ005643 | CRYAB | Small heat shock protein (HSP20) family |
| | gene6864 | 3.43717916 | 0.04992689 | CPIJ004730 | | Chitin-binding domain type 2 |
| | gene9296 | -3.4476402 | 0.04992689 | CPIJ005327 | | |
Supplemental Table 4.2
| DPI | Name | Accession | Gene ID | cor | p-value | Description |
| --- | --- | --- | --- | --- | --- | --- |
| 4dpi | | | nbis-gene-2-utr | 1.00 | 0 | WNV 5' UTR |
| 4dpi | ATRX | CPIJ014697 | gene14505 | 0.69 | 0 | Transcriptional regulator |
| 4dpi | | CPIJ002538 | gene9014 | 0.67 | 0 | Cytochrome p450 |
| 4dpi | | | gene7420 | 0.66 | 0 | |
| 12dpi | | | nbis-gene-2-utr | 1.00 | 0 | WNV 5’ UTR |
| 12dpi | GSTE4 | CPIJ018631 | gene10812 | 0.80 | 0 | Glutathione S-transferase |
| 12dpi | HAO1 | CPIJ013714 | gene13987 | 0.79 | 0.015 | FMN-dependent dehydrogenase |
| 12dpi | METTL20 | CPIJ012802 | gene5405 | 0.77 | 0 | Lysine methyltransferase |
| 12dpi | | | gene870 | 0.75 | 0 | |
| 12dpi | | CPIJ013609 | gene2408 | 0.74 | 0.005 | Myofilin |
| 12dpi | PROX1 | CPIJ004052 | gene10632 | 0.74 | 0.01 | Homeo-prospero domain |
| 12dpi | | | gene3582 | 0.72 | 0 | |
| 12dpi | | | gene9007 | 0.72 | 0 | |
| 12dpi | UROS | CPIJ002599 | gene5208 | 0.72 | 0 | Uroporphyrinogen-III synthase |
| 12dpi | | | gene2263 | 0.71 | 0 | |
| 12dpi | | | gene11056 | 0.71 | 0 | |
| 12dpi | BCAN | CPIJ006816 | gene8287 | 0.70 | 0 | C-Type lectin |
| 12dpi | | | gene10708 | 0.68 | 0 | |
| 12dpi | | CPIJ012060 | gene13467 | 0.67 | 0 | |
| 12dpi | | CPIJ015107 | gene5114 | 0.67 | 0 | Trypsin-like serine protease |
| 12dpi | | CPIJ015715 | gene6473 | 0.67 | 0 | AMP dependent ligase |
| 12dpi | DHDH | CPIJ014580 | gene9019 | 0.67 | 0 | dihydrodiol dehydrogenase |
| 12dpi | | CPIJ009468 | gene5757 | 0.67 | 0 | cytochrome P450 |
| 12dpi | mRpS26 | CPIJ007771 | gene10824 | 0.66 | 0 | Mitochondrial ribosomal protein |
| 12dpi | | CPIJ002678 | gene1254 | 0.66 | 0 | Glutathione S-transferase |
| 12dpi | CHKov1 | CPIJ010305 | gene1164 | 0.65 | 0 | |
| 12dpi | | | gene5412 | 0.65 | 0 | |
| 12dpi | | CPIJ006248 | gene9874 | 0.65 | 0 | Aldo/keto reductase family |

### Slide 19
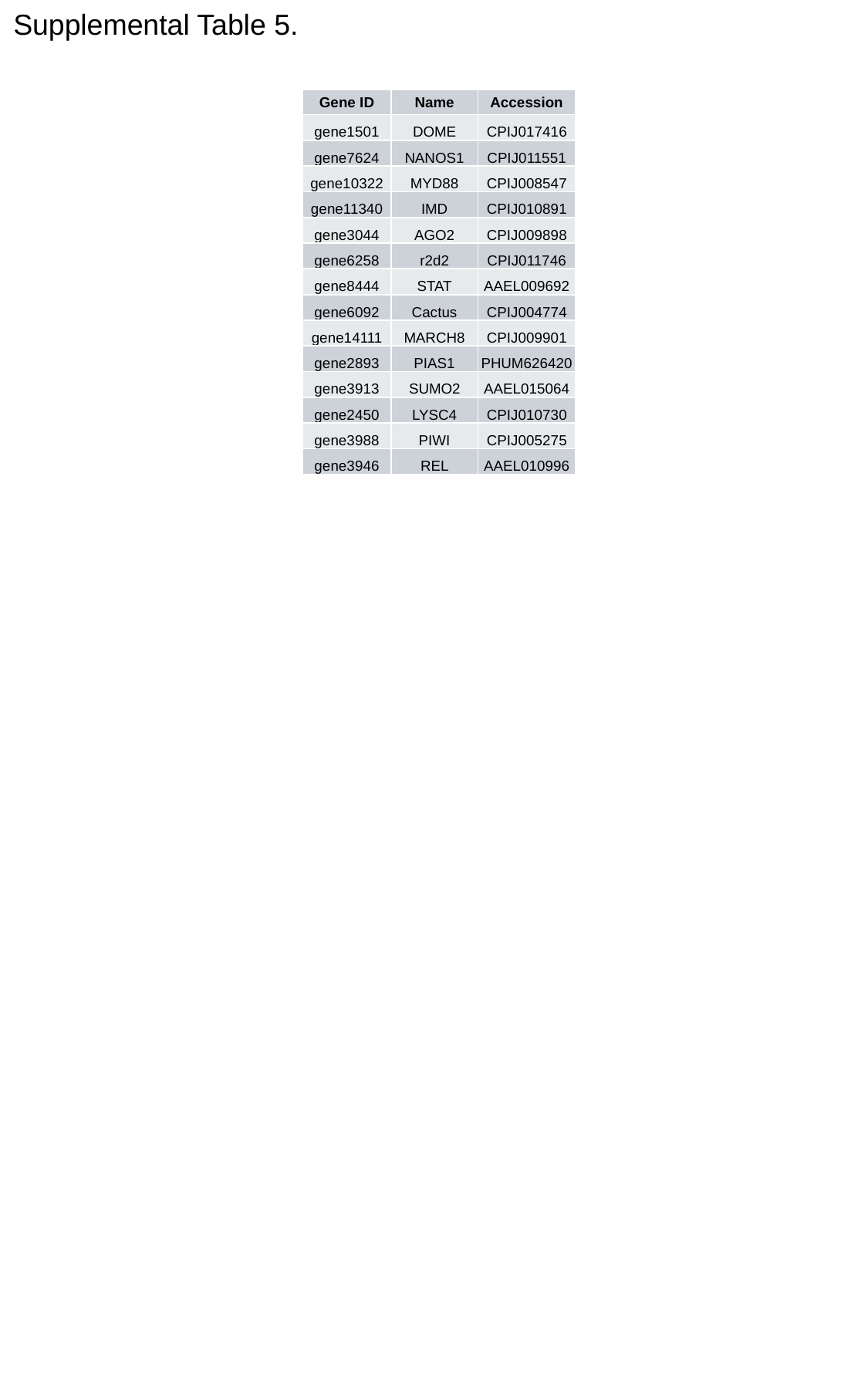

Supplemental Table 5.
| Gene ID | Name | Accession |
| --- | --- | --- |
| gene1501 | DOME | CPIJ017416 |
| gene7624 | NANOS1 | CPIJ011551 |
| gene10322 | MYD88 | CPIJ008547 |
| gene11340 | IMD | CPIJ010891 |
| gene3044 | AGO2 | CPIJ009898 |
| gene6258 | r2d2 | CPIJ011746 |
| gene8444 | STAT | AAEL009692 |
| gene6092 | Cactus | CPIJ004774 |
| gene14111 | MARCH8 | CPIJ009901 |
| gene2893 | PIAS1 | PHUM626420 |
| gene3913 | SUMO2 | AAEL015064 |
| gene2450 | LYSC4 | CPIJ010730 |
| gene3988 | PIWI | CPIJ005275 |
| gene3946 | REL | AAEL010996 |
